# PROTEIN-*O*-MANNOSYLATION BY NON-SEC/TAT SECRETION TRANSLOCONS IN ACTINOBACTERIA

**DOI:** 10.1101/2023.08.23.554508

**Authors:** Hirak Saxena, Rucha Patel, John Kelly, Warren Wakarchuk

**Author notes:** Adress correspondence to Warren Wakarchuk.

## Abstract

Protein-*O*-mannosylation (POM) is a form of *O*-glycosylation that is ubiquitous throughout all domains of life and has been extensively characterized in eukaryotic systems. However, in prokaryotes this process has only been investigated in terms of pathogenicity (in *Mycobacterium tuberculosis*) even though there are many non-pathogenic bacteria that are known to regularly carry out POM. To date, there is no consensus on what benefit POM imparts to the non-pathogenic bacteria that can perform it. Though the generation of a POM deficient mutant of *Corynebacterium glutamicum* – a widely utilized and known mannosylating actinobacteria – this work shows that even closely related actinobacterial GT-39s (the enzymes responsible for the initiation of POM) can potentially have different activities and substrate specificities for targets of POM. Moreover, presented here is evidence that POM does not only occur in a SEC-dependent manner; POM also occurs with TAT and non-SEC secreted substrates in a specific and likely tightly regulated manner. Together these results highlight the need for further biochemical characterization of POM in these and other bacterial species to help elucidate the true nature of its biological functions.

**Importance:** Both the mechanism and overall cellular function of protein-*O*-mannosylation, a ubiquitous subset of *O*-glycosylation, is poorly understood in bacterial systems. In *Mycobacterium tuberculosis* and other pathogenic actinobacteria, numerous secreted virulence factors were identified as mannoproteins, with protein-*O*-mannosylation deficient mutants displaying a less virulent phenotype due to these proteins lacking the modification. However, these findings do not offer any explanations as to why non-pathogenic strains of actinobacteria also perform this modification as in these organisms it is seemingly dispensable. *Corynebacterium glutamicum* is a widely utilized, industrially relevant actinobacteria that also performs protein-*O*-mannosylation. This manuscript describes the utilization of *C. glutamicum* as a Gram-positive recombinant host for the *in vivo* study of actinobacterial protein-*O*-mannosylation and demonstrates the distinct lack of first-hand biochemical data of the process in prokaryotes.

## Introduction

### Actinobacterial Protein O-Mannosylation

POM is an essential and ubiquitous posttranslational modification found throughout all domains of life (1). In the half century since this modification was first described in actinobacteria (2) it has been demonstrated that POM plays a critical role in the virulence of *M. tuberculosis* (3), but no definitive biological context has been attributed to it in the non-pathogenic members of the phylum highlighting a significant lack in the understanding of prokaryotic POM. Most of the information on POM and the enzymes responsible for it – protein*-O*-mannosyltransferases (PMTs) – has been garnered from work done in *S. cerevisiae*, which has contributed significantly to the understanding of this process in eukaryotes. A considerable proportion of the mannosylated eukaryotic proteins identified to date are either secreted or cell membrane associated proteins with well defined functions (1, 4–6). Conversely, only a few of the mannosylated proteins identified in well-known actinobacteria like *M. tuberculosis* (7–9), *C. glutamicum* (10, 11), *Cellulomonas fimi* (12, 13), and *Streptomyces coelicolour* (14, 15) have had functions attributed to them, complicating the elucidation of the biological context of this modification. This is despite there being over 3,000 bacterial species with an annotated PMT – belonging to glycosyltransferase family 39 (GT-39) in the Carbohydrate Active Enzyme (CAZy) database (16).

In eukaryotes, POM is initiated in the lumen of the endoplasmic reticulum (ER) as a protein is being translocated across the membrane in an unfolded SEC-dependent manner. In prokaryotes, POM is thought to occur similarly during extracellular translocation across the plasma membrane; however, many Tat-exported proteins have also been identified to be O-mannosylated, which are folded prior to their export (17–19). Recently, numerous mannosylated cytosolic and non-SEC translocon secreted proteins have also been identified in *C. fimi* and *Cellulomonas flavigena* (20, 21). This contradiction serves to highlight the fact that there is still a significant lack of information on both the mechanisms involved in bacterial protein-O-mannosylation and its overall function.

### POM Requirements

Four mandatory components are necessary for any cell to successfully carry out POM: an activated mannose residue, a mannose-carrying lipid donor, a PMT (GT-39) enzyme, and the target protein to be modified containing the required Ser or Thr residue(s). Though there exists some variation in these components dependent on their parent organism (especially between prokaryotes and eukaryotes) the generalized steps of protein-O-mannosylation are considered to be the same throughout all domains of life (5, 22). There are four generalized steps to protein-O-mannosylation: firstly, the mannosylated lipid donor is synthesized through the action of a GT-2 family glycosyltransferase (known as Ppm1 in actinobacteria), that transfers mannose from the activated donor GDP-α-D-mannose to the phosphorylated prenyl or dolichol donor lipid. Next, the phosphomannose lipid is flipped across the membrane by a still unknown protein to where it can interact with the PMT. The mannose residue is then transferred from the phospholipid carrier to the PMT enzyme, which finally transfers the mannose through an inverting mechanism to the hydroxyl group of Ser or Thr residues on the target protein as it is translocated across a biological membrane (23–28). The most evident difference in protein *O*-mannosylation between eukaryotes and prokaryotes is the cellular location of the modification. As prokaryotes lack the cellular compartments of eukaryotes, protein *O-*mannosylation is thought to occur in the periplasm or on the extracellular face of the plasma membrane (depending on Gram status of the organism) instead of the luminal face of the ER (1, 29, 30).The mono-mannosylated glycoprotein will then undergo further modification by a gamut of distinct enzymes to complete the glycan chain, producing the final glycoprotein product (31). While the enzymes responsible for the subsequent elongation of mannoglycans in actinobacteria is not currently known, it is likely performed by GT-4 family members recognized to have α-1,2-, α-1,3-, and/or α-1,6-mannosyltransferase activity (7 of which are annotated in the *C. glutamicum* ATCC 13032 genome).

### Glycosyltransferase Family 39 (GT-39)

All PMT enzymes identified to date are integral membrane proteins containing several transmembrane domains and contain enough sequence homology to be grouped into GT-39, highlighting the conservation of *O*-mannoslyation (16). In *S. cerevisiae* there are several PMT orthologues belonging to three PMT subfamilies, POMT1, POMT2, and POMT4 (32, 33) which were previously thought to be a form of redundancy; however, it is now known that enzymatic activity requires members of POMT1 and POMT2 to form heterodimers while POMT4 members form homodimers (34, 35). Recently, the first cryo-electron microscopy structure of a GT-39 PMT1-PMT2 heterodimer from *S. cerevisiae* was solved, while only topology reports and hydropathy profiles of several bacterial PMTs have been published (22, 36, 37).

The related architecture of the *S. cerevisiae* GT-39 to GT-66 oligosaccharyl transferases from both prokaryotes (but not including *actinobacteria* as there are no annotated GT-66 enzymes in these organisms) and other eukaryotes implies that PMTs across the domains of life maintain at least three characteristics: a cytosolic N-terminal region and C-terminal region on the opposing side of the membrane (endoplasmic reticulum or plasma membrane), multiple hydrophilic loops on either side of the membrane, with the first luminal/periplasmic loop containing at least part of the PMT catalytic site while carrying the conserved neighbouring residues DE (38–40) aligning with D^55^ and E^56^ in the yeast PMT1. While these two conserved residues are not believed to form part of the PMT active site – as exchange of either does not completely abolish PMT activity – they may be involved in the stabilization of the active site by the coordinating and positioning of the required divalent cations (41). As many studies have focused on characterizing these enzymes in eukaryotic cells, there is precious little first-hand information about the bacterial members of this family (1, 30, 42).

Across species, GT-39s can show a marked degree of variation. The only prokaryotic GT-39s investigated to date originated from *M. tuberculosis* and *C. glutamicum* and show only 25.7% and 26% global sequence similarity, respectively, to the PMT1 of *S. cerevisiae*. Interestingly, these two prokaryotic PMTs share a 55.9% similarity (Table S1), which could be partially explained by their taxonomic relation. Genomic analyses have revealed other putative bacterial GT-39s in other actinobacteria, like *C. fimi* and *C. flavigena* (25.4% and 23.2% similarity to *S. cerevisiae* PMT1, 46% and 46.9% similarity to the *M. tuberculosis* GT-39, and 42% and 40.5% similarity to the *C. glutamicum* GT-39, respectively), including other prokaryotes.

### Lipid Donor and Glycosyltransferase Family 2 (GT-2)

Glycolipid intermediates provide activated mannose to PMTs for POM. There is one specific mannosyl donor in all eukaryotes, and one specific mannosyl donor in all *O*-mannosylating bacteria. These donors are dolichol phosphate β-D-mannose (Dol-P-Man) and polyprenyl monophosphomannose (PPM), respectively (42–44). The interaction of PMTs with the mannosyl donor depends on the presence of an activated mannose moiety anchored to the periplasmic side of the membrane (45, 46) in prokaryotes, or luminal side of the ER for eukaryotes (47, 48).

Prior to the POM reaction a separate synthase enzyme which is anchored into the membrane charges these phosphorylated lipid donors with mannose – using GDP-α-D-mannose (GDP-Man) – on their respective cytoplasmic extension (Maeda & Kinoshita, 2008). These enzymes are polyprenyl monophosphomannose synthases (Ppm1) and belong to the GT-2 family (16). The newly synthesized Dol-P-Man or PPM is then flipped to the opposing side of the membrane, by an unknown protein (5, 49–51) to interact with the PMT and other mannosyltransferase enzymes.

### Transmembrane and Tetratricopeptide Repeat-Containing (TMTC) Mannosyltransferases

Recently a novel class of *O*-Man glycosyltransferases was identified, selectively serving to mono-mannoslyate cadherins and protocadherins (52). The *O*-mannosylglyans on these proteins were reported to not be elongated, suggesting a novel type of *O*-mannosylation in higher eukaryotes (53–56). This enzyme family of transmembrane (TM) and tetratricopeptide (TPR) repeat-containing (TMTC) proteins is composed of four paralogues, TMTC1 – 4, with potentially differing roles (57, 58) and belong to GT-105. While this family is distributed in both eukaryotes and (to a limited degree) prokaryotes, no putative GT-105s currently exist in actinobacteria suggesting that GT-39s are the sole initiating mannosyltransferase in the phylum.

### Investigating Actinobacterial POM in C. glutamicum

*C. glutamicum* is a Gram-positive non-pathogenic, non-sporulating, non-motile rod-shaped bacteria that has been widely utilized in industrial applications such as the production of amino acids, nucleotides, and enzymes (59–61). *C. glutamicum*, *M. tuberculosis*, *C. fimi*, and *C. flavigena* are all members of the Actinomycetales order and have genomes that are high in GC% content. The wide utilization of *C. glutamicum* means there exists an established molecular toolbox for genetic engineering of this organism. In addition, the close genetic resemblance of *C. glutamicum* to *C. fimi* (and common mannosylation machinery) suggests that this organism serves as an ideal candidate for a Gram-positive protein expression/secretion system to investigate actinobacterial mannosylation. This was demonstrated recently, as the *C. glutamicum* platform was shown to accurately mannosylate and localize recombinantly produced heterologous proteins originating from *C. fimi* (21).

A GT-39 deficient mutant of *C. glutamicum* recombinantly producing selected actinobacterial PMTs will allow for the *in vivo* assay of these multipass transmembrane proteins, which are traditionally difficult to biochemically characterize and therefore lacking reports in literature. Using both the native *C. glutamicum* mannoproteome and a natively mannosylated actinobacterial mannoprotein – Celf-3184, expressed alongside each actinobacterial PMT – will show any differences in substrate preference between the related actinobacterial GT-39s, allowing for the continued development of the *C. glutamicum* platform for the investigation of POM in actinobacteria. Most importantly, the impact of different secretion pathways on POM will be investigated using actinobacterial targets, some of which are known to be secreted mannoproteins.

## Results

### Cg_1014 Knockout and Complementation

To adequately assess *in vivo* POM by each recombinantly expressed actinobacterial GT-39, the native *C. glutamicum* GT-39 was inactivated. Homologous recombination was used to knockout the PMT gene in *C. glutamicum* (Cg_1014), taking advantage of the wide host range of the pK18mobsacB (62) suicide vector. Predictive software tools revealed the possibility of regulatory transcriptional elements for neighbouring genes to be contained on the non-coding strand of Cg_1014, specifically, the regions that coded for the N- and C-termini of the PMT enzyme. For this reason, a truncated and inactive knockout construct was designed instead of a seamless knockout (Figure S1). This truncated construct only consisted of the cytoplasmic N-terminal region, the first transmembrane region, and the extracellular C-terminal region. As the active site and conserved residues D^65^ and E^66^ are contained in the first extracellular loop, it was predicted that this construct would effectively abolish POM in *C. glutamicum*. Following homologous recombination, clones were screened and confirmed via colony PCR (Figure S2) for replacement of the native Cg_1014 gene by the truncated and inactive knockout. An amplicon of 2,539 bps indicated a successful knockout generating the ΔCg_1014 strain compared to the 3,880 bps amplicon in ATCC 13032 with intact GT-39. The final, positive clone was further confirmed for the loss of POM by ConA lectin blotting (Figure 2), with any residual ConA reactivity in the POM deficient strain being attributed to mannosylated lipids and/or lipoarabinomannan (LAM) enriched in the membrane fractions via their resistance to proteinase K digestion (Figure S3).

The loss of POM In other actinobacteria has been reported to be non-essential and lacking a discernable phenotype (3, 10). This was confirmed with the GT-39 deficient strain of *C. glutamicum* as there were no significant differences in growth between ATCC 13032 and the ΔCg_1014 mutant (Figure S4). As previous proteomic studies of actinobacteria have identified that many mannoproteins are either membrane-bound or membrane-associated (20, 63), the ΔCg_1014 mutant was further screened for a distinct phenotype with antibiotics targeting either membrane-bound/associated or intracellular components – therefore requiring active or passive transport through the membrane. While no obvious patterns were evident, some differences in antibiotic susceptibility between the two strains were observed (Figure 1). The ΔCg_1014 mutant strain was more susceptible to tetracycline (30 µg), chloramphenicol (30 µg), and tobramycin (10 µg), but less susceptible to novobiocin (30 µg), gentamicin (10 µg), and erythromycin (15 µg) compared to the ATCC 13032 strain. Complementation of the ΔCg_1014 strain with pCGE-31 harbouring the native Cg_1014 gene resulted in antibiotic susceptibilities like the ATCC 13032 strain, except for three of the tested antibiotics. Compared to the ATCC 13032 strain, the complemented mutant was still more susceptible to chloramphenicol (30 µg), tobramycin (10 µg), and erythromycin (15 µg).

**Figure 1:**
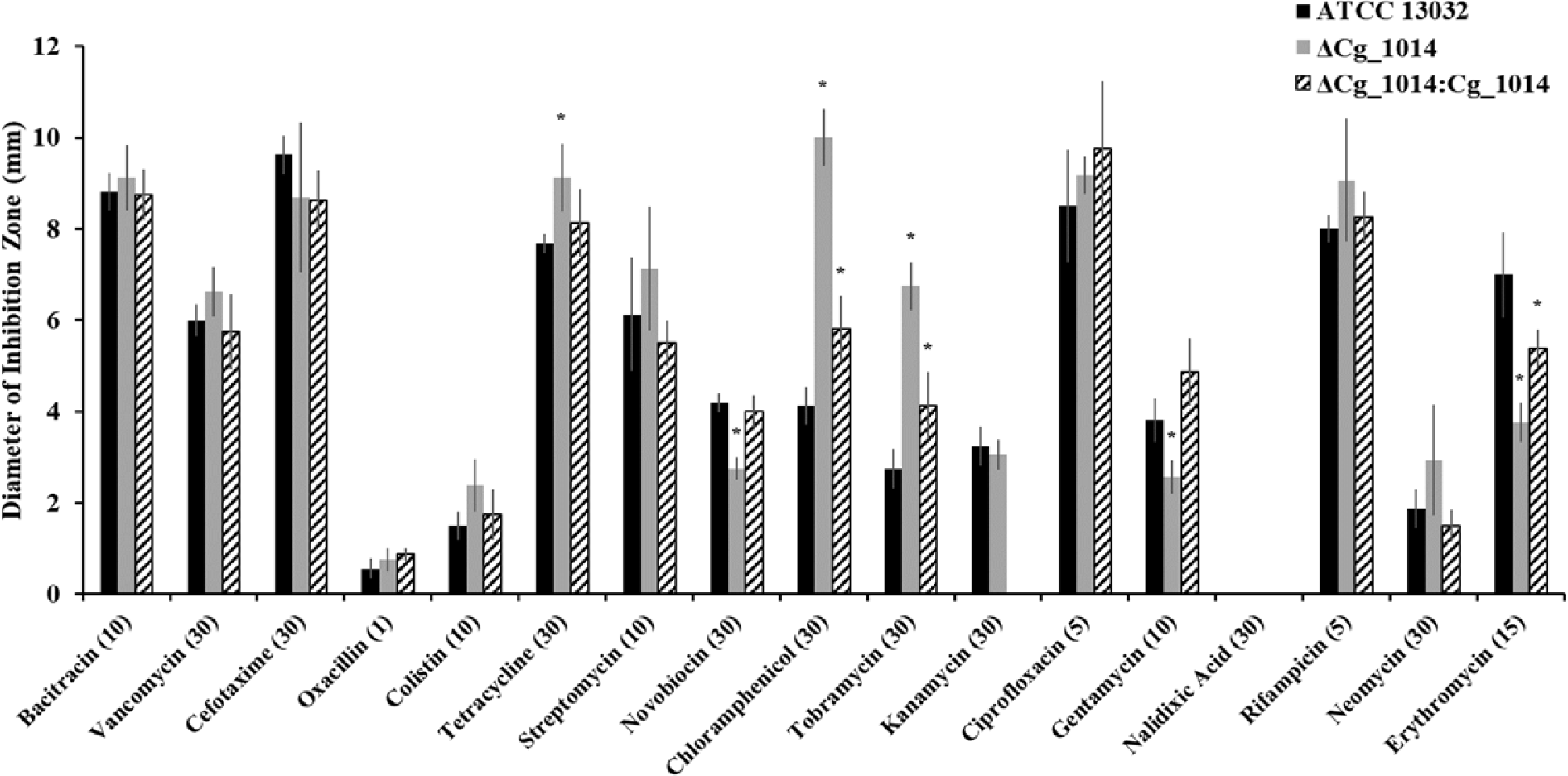
Sensitivity of *C. glutamicum* ΔCg_1014 and ΔCg_1014:Cg_1014 to a range of antibiotics by disc diffusion. Changes to antibiotic sensitivity in the ΔCg_1014 strain were assessed by diameter (in mm) of inhibition zone. The antibiotics having the largest effect on the mutant on ZOI (mm) were tetracycline (30 µg), chloramphenicol (30 µg), tobramycin (10 µg), and erythromycin (15 µg). These antibiotics all inhibit intracellular targets. Complementation of the mutant by Cg_1014 restored sensitivity or resistance to most antibiotics except for chloramphenicol, tobramycin, and erythromycin. These antibiotics target various ribosomal subunits (23S, 30S/50S, and 50S, respectively). Kanamycin resistance of the complemented mutant is conferred by the expression plasmid for antibiotic selection.

**Figure 2:**
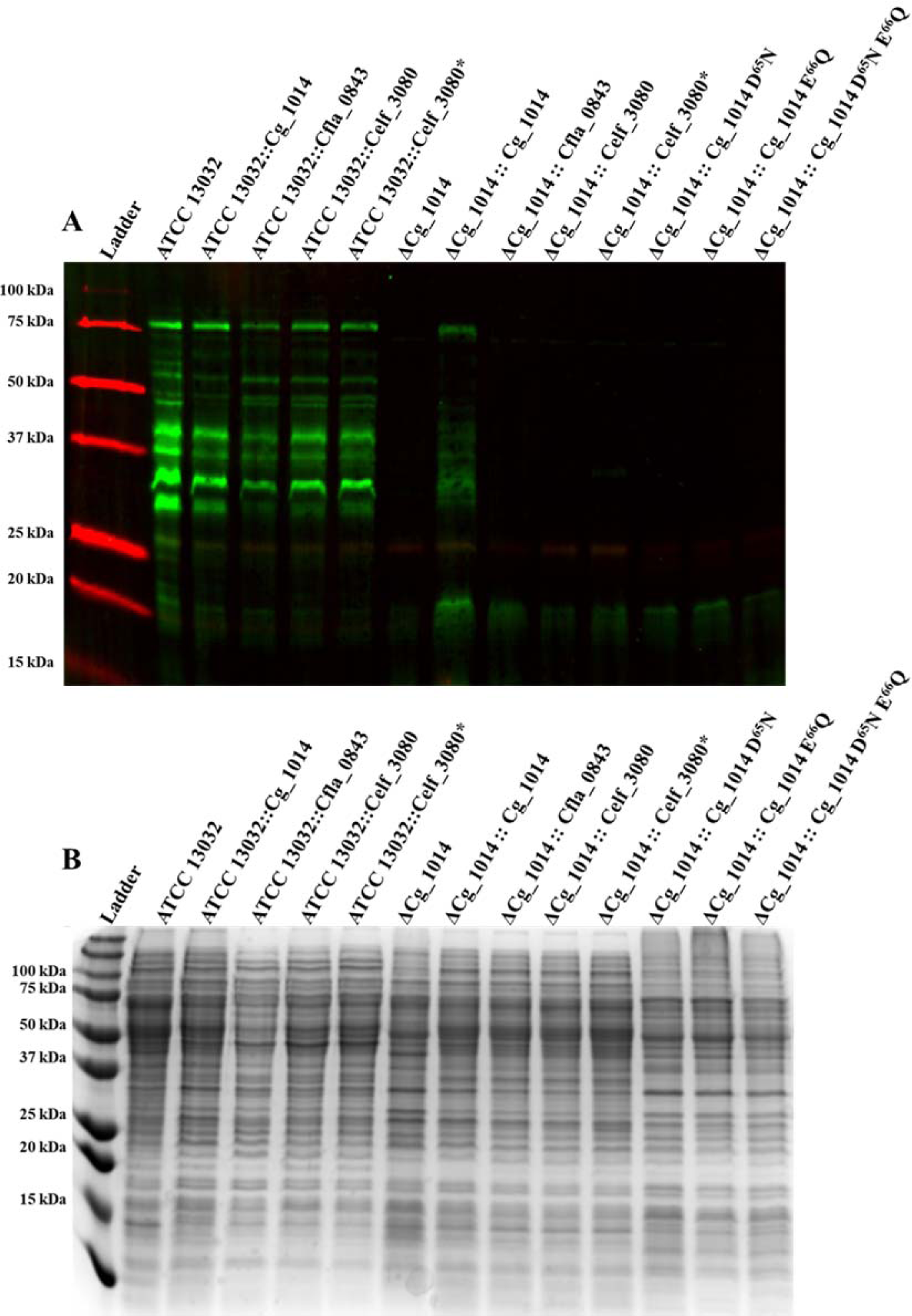
ConA-FITC (green) lectin blot (A) and Coomassie stained 15% SDS-PAGE (B) of *C. glutamicum* ATCC 13032 and ΔCg_1014 membrane fractions expressing recombinant actinobacterial GT-39s and Cg_1014 SDM constructs. Overexpression of the actinobacterial GT-39s in *C. glutamicum* ATCC 13032 resulted in no changes to the native membrane protein mannoproteome as detected by Concanavalin A-FITC lectin conjugate (A, green). Western blotting using anti-HIS_6_ antibodies and fluorescent Ni-NTA conjugates did not detect any proteins indicative of recombinant GT-39s. Lack of POM in the ΔCg_1014 strain was complemented by expression of the *C. glutamicum* GT-39 Cg_1014, with minimal complementation by the *C. fimi* GT-39 Celf_3080, and no complementation was evident with the *C. flavigena* GT-39 Cfla_0843. Substitution of the conserved residues D^65^ and E^66^ to N^65^ and Q^66^, respectively, confirmed their requirement for catalytic activity. An asterisk (*) denotes the *C. fimi* GT-39 (Celf_3080) codon optimized for expression in *C. glutamicum*. Coomassie stained 15% SDS-PAGE as loading control (B). Molecular weight standards are the Bio-Rad All Blue ladder.

The traditional biochemical characterization of multipass transmembrane proteins is notoriously difficult and problematic (64, 65). Detection and recovery of actinobacterial GT-39s produced recombinantly in both the ATCC 13032 and ΔCg_1014 strain was not possible (data not shown). For this reason, the *O*-mannosylation of the native *C. glutamicum* mannoproteome was initially used to assess the degree of POM in the complemented ΔCg_1014 strains. Only the ΔCg_1014 strain complemented with the *C. glutamicum* GT-39 showed notable recapitulation of mannosylation, with minimal complementation by the *C. fimi* GT-39, Celf_3080 (Figure 2). Lack of complementation in the ΔCg_1014 strain with SDM mutants of the *C. glutamicum* GT-39 – mutating conserved residues D^65^N and E^66^Q – suggests that these residues are necessary for enzymatic activity, but not necessarily catalysis (Figure 2). Like the *S. cerevisiae* and *S. coelicolor* GT-39s, it is likely that these residues assist in the coordination of an active site divalent Mn2^+^ cation – based on their structural similarity to the oligosaccharyltransferase from *Campylobacter lari*, PglB, where activity is metal dependent (36, 66, 67).

### In vivo O-Mannosylation using an Actinobacterial Target Mannoprotein

As the native *C. glutamicum* mannoproteome proved to be a poor target for the *C. fimi* and *C. flavigena* GT-39s, a selected target actinobacterial mannoprotein – Celf_3184, known to be accurately mannosylated in the ATCC 13032 strain (21) – was expressed alongside each actinobacterial GT-39 in a synthetic operon to better assay the *in vivo* mannosylation of these PMTs. Interestingly, differential mannosylation of Celf_3184 was detected when co-expressed with the different actinobacterial GT-39 complementation constructs (Figure 3). Perhaps unsurprisingly, LC-MS results confirmed that the culture supernatant recovered Celf_3184 co-expressed with the *C. fimi* GT-39 in the ΔCg_1014 strain was more uniformly modified by hexoses (Figure 4A) than when produced in the ATCC 13032 strain, which has been previously reported. When produced in the ATCC 13032 strain (or the *C. glutamicum* GT-39 complementation construct) the range of modifications to Celf_3184 typically falls between 29 – 37 hexoses (21), but when co-expressed with the *C. fimi* GT-39 this range is much narrower at 31 – 35 hexoses (Figure 4A). Celf_3184 co-expressed with the *C. glutamicum* GT-39 in the ΔCg_1014 strain showed a similar degree of modification (data not shown) to Celf_3184 produced in the ATCC 13032 strain (21). Conversely, the *C. flavigena* GT-39 does not appear to accept any of the *C. glutamicum* native mannoproteome or Celf_3184 as POM targets which could suggest a very stringent substrate specificity for this GT-39.

**Figure 3:**
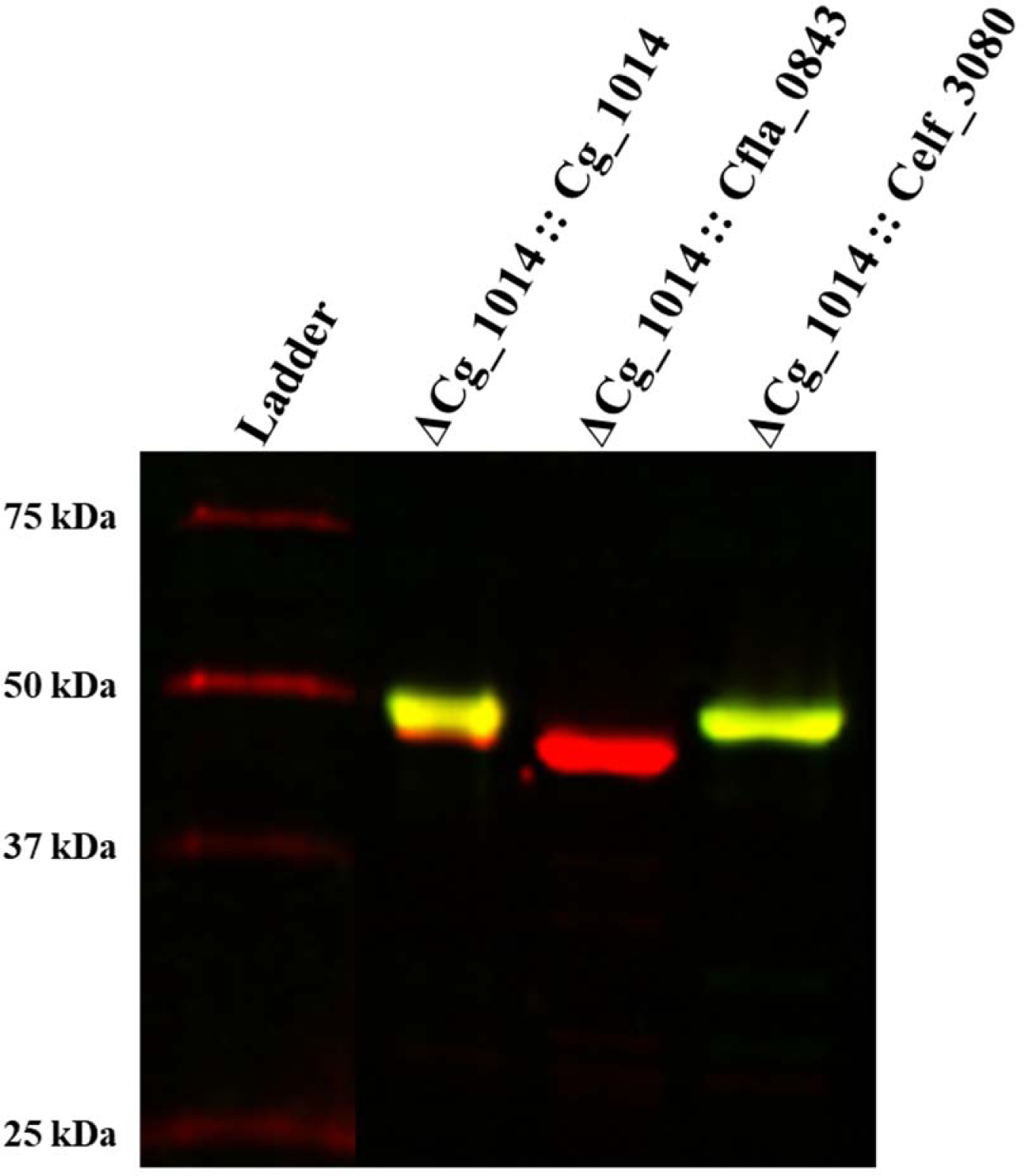
ConA-FITC (green) and Anti-HIS-Alexa647 (red) blot of recombinant Celf_3184 recovered from spent media of *C. glutamicum* ΔCg_1014 expressing actinobacterial GT-39s. As direct detection of recombinant GT-39s was not possible, Celf_3184 was used as an in vivo readout of POM activity. When co-expressed with each actinobacterial GT-39 in the ΔCg_1014 strain both the *C. glutamicum* and *C. fimi* GT-39s accepted the known mannoprotein as a substrate, with the *C. fimi* GT-39 producing a more homogenously mannosylated product. The GT-39 of *C. flavigena* did not glycosylate this substrate, but it was correctly exported out of the cell. Molecular weight standards are the Bio-Rad All Blue ladder.

**Figure 4:**
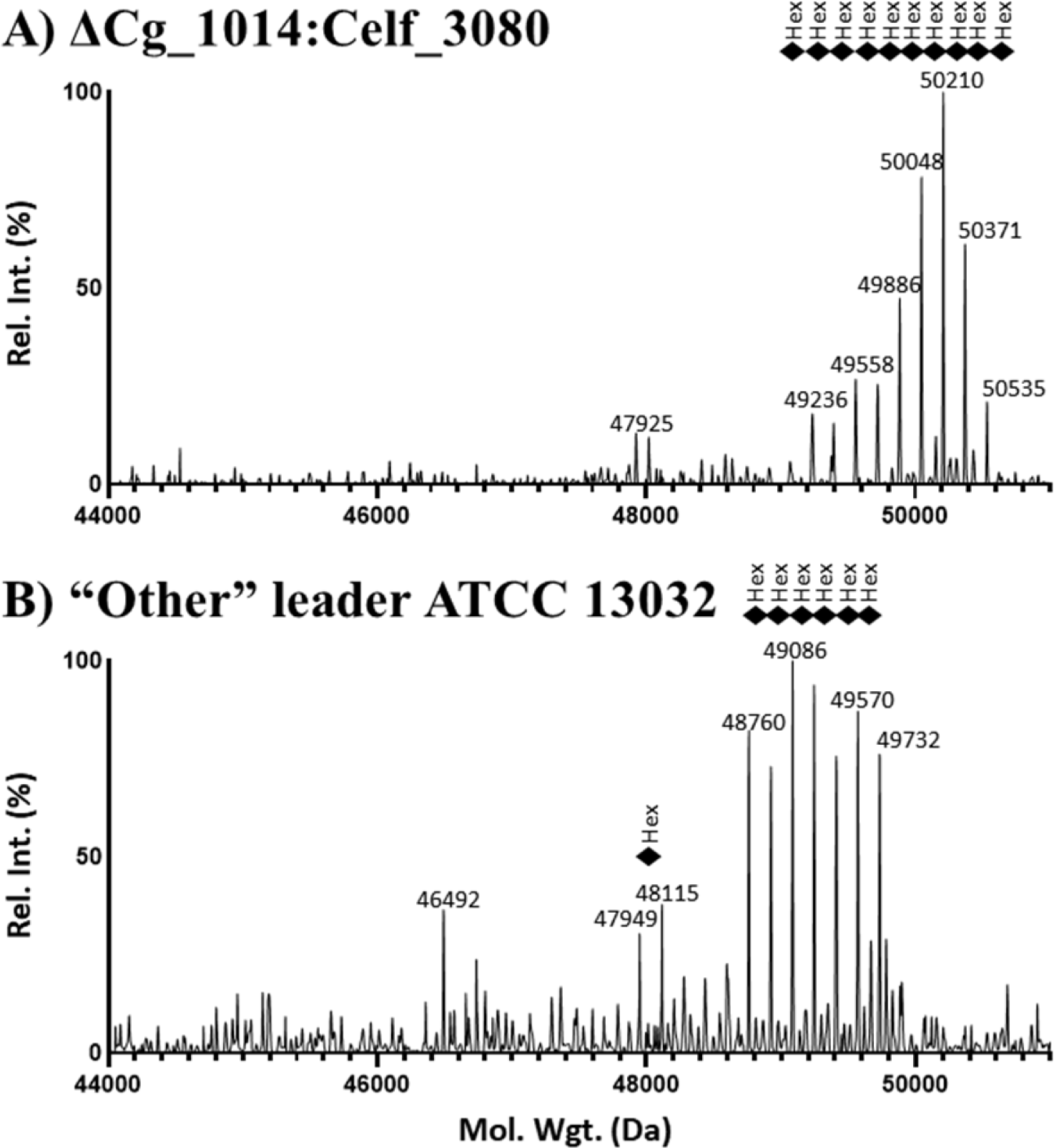
Intact mass LC-MS analysis of Celf_3184 expressed in (A) ΔCg_1014:Celf_3080 and (B) ATCC 13032 using “Other” leader sequence from Celf_2022. Both proteins were recovered from the spent culture media. The calculated mass (with signal peptide removed and no hexose modifications) of Celf_3184 is 44,880 Da. When co-expressed with the *C. fimi* GT-39 Celf_3080 in the ΔCg_1014 strain, the observed mass profile (A) suggests this protein is modified with between 31 – 35 hexoses, which is much more homogenous than the modification seen on native Celf_3184 when produced in ATCC 13032. When expressed in ATCC 13032 and with its native TAT leader replaced by the “Other” leader sequence of Celf_2022, the produced Celf_3184 has an observed mass profile (B) suggesting the addition of 24 – 30 hexoses; fewer hexose modifications than when native Celf_3184 is produced in ATCC 13032.

### Secretion and POM Utilizing a Translocon Other than SEC and TAT

Originally, predictive tools were unable to identify an *N*-terminal secretion signal associated with Celf_2022 even though it was first identified as a mannoprotein lacing a predicted leader sequence (a predicted cyclophilin type peptidyl-prolyl cis-trans isomerase) in the spent media of *C. fimi* cultures during the proteomic analysis of the secretome of *C. fimi* (20). For this reason, Celf_2022 was initially chosen as a possibly cytoplasmic mannoprotein target for heterologous expression in *C. glutamicum*. The previous validation of *C. glutamicum* for the accurate mannosylation of recombinant actinobacterial mannoproteins showed that the recombinant host is capable of producing, exporting, and mannosylating Celf_2022 similarly to when it is produced in its native organism *C. fimi*. Further, it was determined that Celf_2022 is actually a highly efficiently secreted mannoprotein, with the identified glycopeptide at the N-terminus of the mature polypeptide (21). While bioinformatic tools are often capable of accurately identifying and classifying signal peptides in addition to predicting protein localization, the potential for misclassification must be noted as in the case of Celf_2022. In contrast to the predicted cytoplasmic localization, significant amounts of this protein were found to be secreted into the culture media when produced in *C. glutamicum* (21).

Subsequent iterations of the SignalP algorithm now classify the leader sequence of this protein (likelihood of 0.998 using SignalP 6.0) as likely utilizing a pathway other than SEC or TAT, leading to the the Celf_2022 leader sequence being classified as “Other”. To confirm this prediction, the “Other” leader sequence of Celf_2022 was exchanged with the SEC leader of Celf_1230 and the TAT leader of Celf_3184. As a functional assay for this enzyme has yet to be developed, POM was used to assess which translocon could possibly be utilized by this protein during native secretion. These two translocons are both involved in the process of actinobacterial POM but differ mechanistically. Proteins that are secreted through the SEC translocon are exported in an unstructured or linear manner, while the TAT translocon secretes proteins that are in a fully folded state. A matching POM profile between one, or both leader swapped Celf_2022 constructs and the protein with its native “Other” leader would then be a strong indicator of which translocon was being utilized by the “Other” leader sequence.

### Celf_2022

Celf_2022 is mannosylated (and exported extracellularly) with its native “Other” leader sequence in ATCC 13032, but replacement by either a classical SEC leader (from Celf_1230) or a classical TAT leader (from Celf_3184) completely abolished mannosylation in the secreted recombinant protein that was recovered from the spent culture media (Figure 5A and D). The low levels of recombinant Celf_2022 within the cells (Figure S5A and B) confirmed that the protein observed in the unconcentrated culture media (Figure S5C and D) was not due to cell lysis. To confirm POM of native Celf_2022 was carried out by the *C. glutamicum* GT-39 alone, all Celf_2022 constructs were expressed in the ΔCg_1014 strain where no mannosylation was detected on any protein regardless of the secretion leader utilized (Figures S6A and D, S7). These results support the predictive analysis of SignalP 6.0, suggesting that the native Celf_2022 is likely secreted by a translocon independent of both SEC and TAT as only the “Other” leader results in mature Celf_2022 with a POM profile matching that of the original protein identified from the secretomic analysis of *C. fimi* (20).

**Figure 5:**
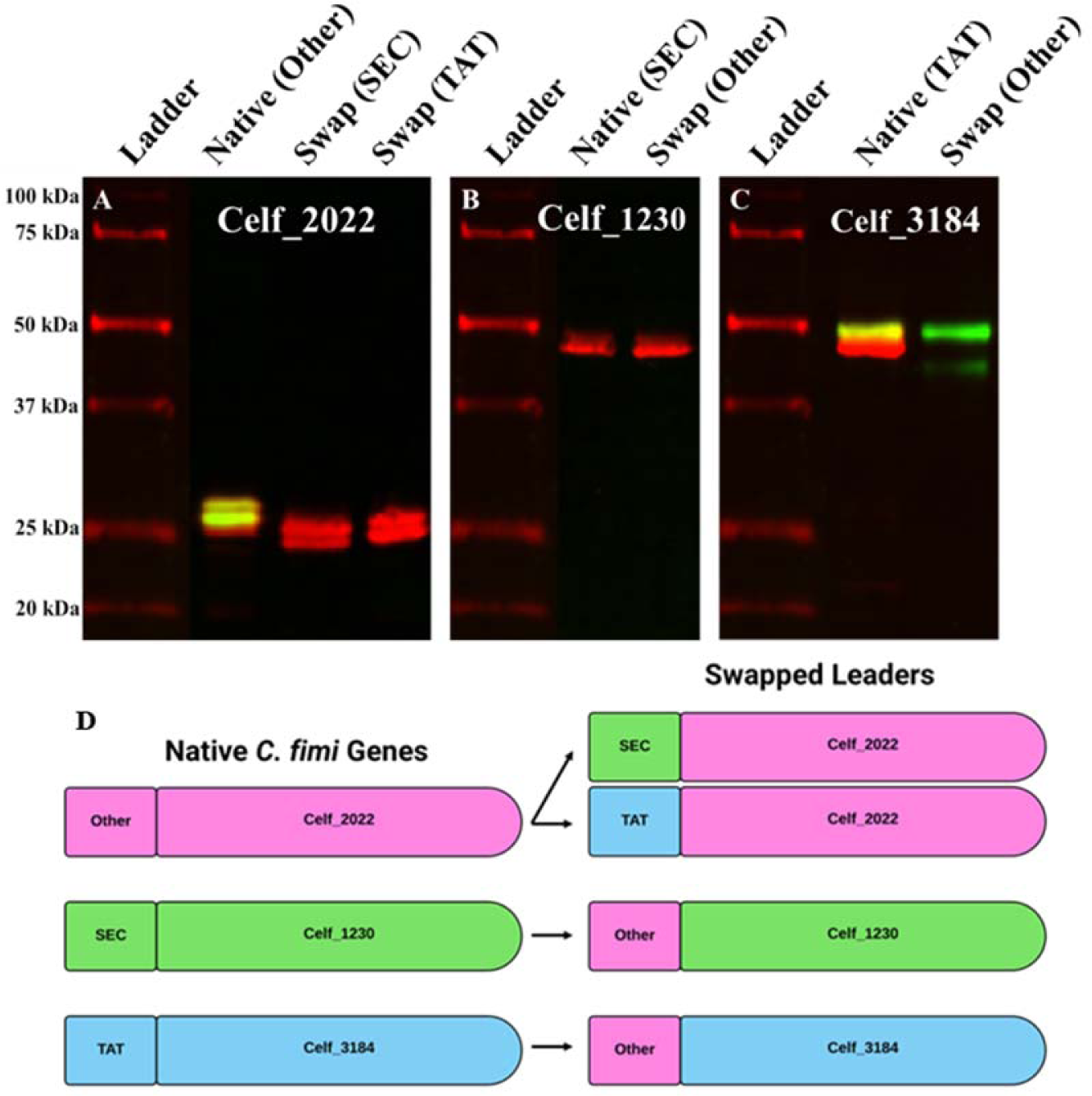
ConA-FITC (green) and Anti-HIS-Alexa647 (red) blot of spent culture media enriched recombinant Celf_2022 expressed with “Other”, SEC, and TAT leaders (A), Celf_1230 expressed with SEC and “Other” leaders (B), and Celf_3184 expressed with TAT and “Other” leaders (C) produced in *C. glutamicum* ATCC 13032. Schematic of swapped leader constructs (D). When lead by its native “Other” leader sequence, Celf_2022 is both secreted and mannosylated in *C. glutamicum* ATCC 13032 (A). Replacement of this “Other” leader sequence by a SEC or TAT signal peptide retains secretion but abolishes mannosylation. Replacement of the SEC signal peptide of Celf_1230 (a non-mannosylated protein) by the “Other” leader sequence results in no change to secretion or mannosylation pattern (B). Replacement of the TAT signal peptide of Celf_3184 by the “Other” leader sequence results in a more homogenous glycoform of protein enriched from the culture media (C). Schematic diagram showing swapped leader sequences between recombinantly produced *C. fimi* test proteins (D). Molecular weight standards are the Bio-Rad All Blue ladder.

### Choice of Secretion Pathway Impacts O-Mannosylation Profile

As the “Other” leader still results in efficient mannosylation of Celf_2022 and was shown to be adjacent to both the SEC and TAT translocons, the effects of this leader on the SEC secreted Celf_1230 and TAT secreted Celf_3184 were investigated. While both proteins were also identified during the secretomic analysis of *C. fimi* (20), only Celf_3184 is known to be a mannoprotein. Like the previous experiment, the effects of exchanging the native leader sequences of these proteins with the “Other” leader sequence of Celf_2022 were assessed by comparing the POM profiles of the leader swapped constructs to the proteins with their native leader. Again, the relative abundance of recombinant proteins in the cytoplasm (Figure S5A and B) compared to the unconcentrated spent culture media (Figure S5C and D) confirmed their presence was not due to cell lysis and constructs were expressed in the ΔCg_1014 strain to confirm POM was carried out by the *C. glutamicum* GT-39 (Figures S6B, C, and D, S7).

### Celf_1230

Celf_1230 is a uniquely thermostable glycoside hydrolase family 6 (GH-6) enzyme from *C. fimi* not identified as a mannoprotein (20, 68) with ConA lectin blotting of the recombinantly produced enzyme in *C. glutamicum* confirming this (Figure 5B). Swapping the native SEC leader of this enzyme with the “Other” leader did not change the mannosylation status of the secreted material (Figures 5B and D, S5C and D). However, relative protein abundance of the “Other” swapped Celf_1230 was greater in both the cytoplasm and unconcentrated spent media fractions (Figure S5), possibly indicative of either greater stability/proteolytic resistance of the non-native fusion.

### Celf_3184

Celf_3184 is another *C. fimi* GH-6 enzyme identified previously (20), different from Celf_1230 in that the former is a highly mannosylated and TAT secreted enzyme. As this protein is an excellent example of a mannoprotein outside of the classical dogma of SEC-dependent POM in actinobacteria it was originally selected for assessing *C. glutamicum* for its use in producing accurately mannosylated recombinant actinobacterial proteins. When produced in *C. glutamicum*, Celf_3184 – with its classical TAT type leader sequence – is mannosylated with between 29 – 37 hexoses (21) which is congruent to observations seen from the same protein produced natively in *C. fimi* (20, 21). However, replacement by the “Other” leader sequence from Celf_2022 resulted in less secreted protein with a more uniform POM profile (Figure S5C and D). Material recovered from the spent culture media (Figure 5C and D) was determined to have 24 – 30 overall hexoses by LC-MS/MS (Figures 4B). The cytoplasmic fraction also showed a lower abundance of Celf_3184 with the “Other” leader sequence (Figure S5A and B) possibly indicative of either decreased stability/proteolytic resistance of the non-native fusion. This appears to be the first report of differential POM on the same target protein in actinobacteria based on the secretion leader sequence utilized.

## Discussion

### C. glutamicum GT-39 is Dispensable

Inactivation and complementation of the gene encoding the GT-39 of *C. glutamicum* has been reported previously (10), however only the presence or absence of mannosylation on secreted proteins was assayed; the more abundant membrane associated or even cytoplasmic mannoproteins were not considered. In addition, no phenotype or growth characteristics of this knockout have been reported to date. For this purpose, the inactivated truncation mutant which maintains up- and downstream effectors (Figure S1B) for Cg_1013 – a hypothetical protein with unknown function – and Cg_1015 – a uroporphyrin-III C/tetrapyrrole (corrin/porphyrin) methyltransferase (69) – was designed to be a more refined approach to generating the mutant required for the investigation of POM in *C. glutamicum*.

Initially, the presence of some ConA reactive bands and smears on lectin blots (Figure 2) led to the assumption of incomplete inactivation of Cg_1014, but this reactivity can be attributed to mannosylated lipids and/or LAM enriched during the isolation of membrane proteins (Figure S3). It has been reported that POM deficient *M. tuberculosis* exhibits enhanced release of LAM (70), which was also observed with the ΔCg_1014 mutant of *C. glutamicum*. While the ConA reactive smears in Figure S3B are indicative of LAM in the samples, it should be noted that there are still some weak but distinct ConA reactive bands in ATCC 13032 following proteinase K digestion. If these bands are in fact proteinaceous in nature, they likely correspond to mannoproteins (or mannose containing glycopeptides) not digested by proteinase K as POM is known to confer proteolytic resistance to some proteins (71, 72). In *M. tuberculosis,* LAM is mannosylated by a mannosyltransferase known as pimB (Rv0557), belonging to GT-4 (73, 74) – a family with no annotated protein-*O*-mannosylation functionality (EC 2.4.1.109). Orthologues of the *M. tuberculosis* GT-4 exist in *C. glutamicum* and will be discussed later. Much like in *M. tuberculosis* (3), inactivation of the *C. glutamicum* GT-39 results in no discernable growth phenotype (Figure S4) exemplifying the seemingly dispensable nature of this modification in non-pathogenic actinobacteria, at least under laboratory conditions.

While microscopy showed no obvious morphological differences between the strains (data not shown), differences in antibiotic susceptibility between ATCC 13032 and ΔCg_1014 were seen with tetracycline, novobiocin, chloramphenicol, tobramycin, gentamicin, and erythromycin (Figure 1) – antibiotics primarily targeting Gram positives and all with intracellular targets. These antibiotics all require transport (passive or active) across the cell membrane, so these results suggest that POM does affect either permeability or specific uptake mechanisms at least in the cases of tetracycline, chloramphenicol, and tobramycin. Surprisingly, lack of POM increased the resistance of the ΔCg_1014 strain to novobiocin, gentamicin, and erythromycin. These antibiotics can be taken up by active transport – using ABC transporters, efflux pumps, or even porins (75) – which have been characterized in *C. glutamicum* (76) – suggesting that possibility that key enzymes along these uptake routes may require POM for functionality. Hexose modified ABC transporters have been reported in *M. tuberculosis* (77). In analogy with the non-functional PMT mutants made in *S. coelicolor* (78), only mild increases to antibiotic susceptibility are seen with vancomycin and β-lactams, which were not observed with the ΔCg_1014 strain. This suggests that in *C. glutamicum*, major cell wall effects are not a phenomenon associated with a lack of protein mannosylation.

Complementation of the mutant by Cg_1014 mostly restored antibiotic susceptibilities to that of the ATCC 13032 strain, with three exceptions. Compared to the wild-type strain, partial complementation of antibiotic susceptibility to chloramphenicol, tobramycin, and erythromycin was observed when the ΔCg_1014 strain was complemented with Cg_1014 – these antibiotics all targeting various ribosomal subunits: 23S, 30S/50S, and 50S, respectively (79–81). This is likely related to POM in the complemented mutant not returning to levels observed in the ATCC 13032 strain (Figure 2). Taken together, these data implicate POM in affecting the interaction between some ribosomal subunits and the antibiotics targeting them. While investigations into the cytoplasmic mannoproteome of actinobacteria is lacking, proteomic evidence of ribosomal subunits in *M. tuberculosis* carrying hexose modifications has been reported (77).

The wide reach of POM – in terms of the biological functions of the proteins carrying the modification – makes it difficult to clearly assign any singular cellular role to it. The abundant nature of POM further complicates the answer to the fundamental question: what biological function does protein-*O*-mannosylation have in these non-pathogenic actinobacterial species?

### Differential Complementation of ΔCg_1014 by Actinobacterial GT-39s

Verification of the actinobacterial GT-39 expression constructs was first attempted using the ATCC 13032 strain. However, no anti-HIS^6^ reactive bands of corresponding molecular weight were identified in the membrane or intermediate fractions of any construct (data not shown) and all *C. glutamicum* expression strains were subsequently confirmed to harbour the correct vector and insert by plasmid rescue into *E. coli*. As a proxy for direct detection, changes to the ATCC 13032 native mannoproteome during the expression of each actinobacterial GT-39 was also assessed (Figure 2); however, no noticeable changes were observed. Integral membrane proteins can be notoriously difficult to detect, enrich, and biochemically characterize (64, 65) as they are often presented at very low levels and lose their native activity and the conformation required for enzymatic activity when removed from biological membranes (82–84). As the *C. glutamicum* vector pCGE-31 typically produces recombinant proteins to a high degree in the organism (Figure S9), it was thought that the majority of the recombinant GT-39s produced in each construct were misfolded and then rapidly degraded. In *E. coli*, the ATP-dependent zinc metalloprotease FtsH degrades misfolded membrane proteins (85); the *C. glutamicum* FtsH homologue (Uniprot A0A1Q6BR68) likely performing a similar function and explaining the lack of detection of even insoluble, misfolded recombinant GT-39s. In *S. coelicolor*, evidence has been presented that even very conservative single amino acid substitutions to the native GT-39 sequence may cause misfolding and rapid degradation leading to both lack activity and detection by Western blotting (86). It is therefore not unreasonable to think that either the *N-* or *C*-terminal HIS^6^ tagging of the actinobacterial GT-39s utilized here could have a similar effect.

Following this reasoning, we sought to observe if enzymatic activity was observable using the ΔCg_1014 strain instead, where any low-level recombinant GT-39 activity would not have to compete with the natively produced enzyme for mannoprotein substrates. In the ΔCg_1014 strain, the *C. glutamicum* GT-39 complemented POM to a degree, but mannosylation levels did not return to that of the ATCC 13032 strain (Figure 2). As enzymatic activity is present in this complemented strain with no recombinant protein able to be detected, it is likely that at least a minimal amount of the *C. glutamicum* GT-39 enzyme is produced in an active form and localized correctly. Knowing this, we attempted to confirm the necessity of the conserved DE motif for enzymatic activity in the first extracellular loop of the *C. glutamicum* GT-39 with site directed mutants. Mutation of either D^65^N or E^66^Q totally abolishes the enzymatic activity of the *C. glutamicum* GT-39. This finding does not prove that the residues are required for catalysis, instead highlighting their importance as conserved residues within the GT-39 family as the DE motif has been implicated in the stabilization of the GT-39 active site by coordinating the required divalent cations (41). Minimal complementation of select mannoproteins was observed with the *C. fimi* GT-39 (native and codon optimized for *C. glutamicum* expression), again suggesting that perhaps a minimal amount of recombinant enzyme was functional. However, no obvious complementation by the *C. flavigena* GT-39 was evident. While this could suggest that these closely related GT-39s may display high specificity for their native targets of POM, the possibility of no active *C. flavigena* GT-39 being produced could not be ruled out. For this reason, we decided to utilize the highly mannosylated protein Celf_3184 (Cel6A) from *C. fimi* as a recombinant target for the *in vivo* readout of POM activity by heterologous GT-39s in *C. glutamicum*.

### In vivo Mannosylation of Celf_3184 by Actinobacterial GT-39s

The *C. fimi* GH-6, Celf_3184, was chosen as the preliminary target mannoprotein for the *O*-mannosylation operons as it has previously been shown to be a suitable target for recombinant expression and POM in *C. glutamicum* ATCC 13032. As this protein is heavily mannosylated in both its native organism, *C. fimi*, and the recombinant host *C. glutamicum* (21), this target mannoprotein was utilized to assess and compare the *in vivo* activities of the different actinobacterial GT-39s expressed in *C. glutamicum* (21). When expressed in conjunction with the different actinobacterial GT-39s differences in mannosylation activity can be seen, but also possible substrate preferences between these closely related enzymes (Figure 3). Logically, the *C. fimi* GT-39 (with native codon usage) readily accepts the *C. fimi* GH-6 and mannosylates it much more uniformly than the *C. glutamicum* GT-39 (Figure 4A), where different glycoforms are all secreted into the culture media. GT-39s only catalyze the initial attachment of mannose to S/T residues and do not contribute to their elongation and as such, the increased degree of modification observed with the *C. fimi* GT-39 cannot be solely attributed to its presence. However, as previously reported most of the *O*-mannose glycans on Celf_3184 (produced recombinantly in ATCC 13032) are disaccharides and not further polymerized (21). As the initial attachment of mannose to S/T residues is likely the limiting step in POM these results show that the *C. fimi* GT-39 recognizes and acts on a greater number of potential *O*-mannosylation sites contained in Celf_3184 than the *C. glutamicum* GT-39. However, the *C. flavigena* GT-39 does not appear to accept this substrate as a target for POM. Again, without a known *C. flavigena* target protein the total lack of active GT-39 production cannot yet be ruled out for this strain. However, *C. flavigena* harbours its own orthologue of the *C. fimi* TAT secreted GH-6 (Cfla_2913) and interestingly, the only significant structural difference between these two related enzymes is in the linker region between the CBM2a and GH6 domains which contains the identified glycopeptide of Celf_3184 (21). The linker region in Cfla_2913 is significantly larger than that in Celf_3184, potentially moving the region homologous to the Celf_3184 glycopeptide (Figure S8). As the N-terminus of Cfla_2913 contains TAT motifs similar to those found in Celf_3184, it is likely also secreted via the TAT translocon where respective GT-39s should be interacting with these proteins in a fully folded state. While Cfla_2913 has yet to be identified as a mannoprotein, if it is natively mannosylated similarly to its *C. fimi* counterpart then these preliminary results imply that there are notable differences in both the structure and spatial functionality of closely related actinobacterial GT-39s. The further dissection of this observation and actinobacterial POM in general requires additional mannoprotein targets from each species be incorporated into the mannosylation operons. Comparing these mannosylation profiles could potentially allow for the identification of a minimum sequence requirement for actinobacterial POM.

### “Other” Leader of Celf_2022 Does Not Utilize SEC or TAT Translocons

To further investigate the potential of Celf_2022 utilizing a translocon other than SEC or TAT, the “Other” leader sequence of the protein was exchanged with the leader sequences from both Celf_1230 (a non-mannosylated SEC secreted protein) and Celf_3184 (a heavily mannosylated TAT secreted protein). While all Celf_2022 constructs were both secreted from *C. glutamicum* ATCC 13032 and enriched from spent culture media, only the native Celf_2022 with “Other” leader sequence was ConA reactive (Figure 5A and D). As the SEC and TAT translocons are known to secrete proteins in unfolded and folded states, respectively, the loss of mannosylation by directing Celf_2022 through these two translocons provides evidence that the context of the glycosylation site is determined by the direction provided by the leader sequence – especially as the identified glycopeptide is totally separate from the native leader sequence which is processed following secretion. Celf_2022 is still an uncharacterized protein with a positively charged *N*-terminus followed by a transmembrane helix that spans residues 33 – 54, the same location where previous iterations of the SignalP algorithm predicted processing of the signal peptide and where the *N*-terminus of the secreted protein was determined to be (21). While these features make the protein appear similar to a single-pass transmembrane protein that likely follows the positive-inside rule, proteomic evidence suggests that this protein is fully secreted instead of membrane associated (20). Regardless of the true location of the mature protein outside of the cell, membrane insertion and secretion both require a manner of translocation through the bilayer and the evidence presented here suggests that in the case of the “Other” leader of Celf_2022 the translocon that is used is neither SEC nor TAT. Finally, To confirm that the mannosylation of native Celf_2022 was not carried out by a separate class of mannosyltransferases (like in the case of LAM), these constructs were also expressed in the ΔCg_1014 mutant where POM was not evident on any recombinantly produced protein (Figure S6A, B, and C). The observed mass differences in both mannosylated and non-mannosylated Celf_2022 is the result of differential *N-*terminal truncation products (21).

Definitive confirmation that Celf_2022 is secreted in a TAT independent manner could be accomplished with the knockout of the *C. glutamicum* TatABC translocase as previously reported (87), but showing the protein is secreted in a totally SEC-independent manner may not be possible as the total knockout of the SEC translocase is known to be lethal in some bacterial species (88) and efforts to do so have yet to be reported in *C. glutamicum*.

### Secretion Translocons Can Modulate POM

To investigate the effects on POM by the “Other” leader and the currently unidentified translocon utilized by it, both Celf_1230 and Celf_3184 had their native leader sequences replaced with the “Other” leader sequence of Celf_2022. Celf_1230 was not detected as a mannoprotein in the proteomic analysis of secreted *C. fimi* proteins (20); its mannosylation profile does not change when the native SEC leader is replaced by the leader sequence of Celf_2022 (Figure 5B and D). This shows that utilizing the “Other” leader and translocon does not result in the aberrant mannosylation of natively non-mannosylated proteins and further reinforces the sole GT-39 of *C. glutamicum* being responsible for all POM.

When the native TAT leader of Celf_3184 is replaced by the Celf_2022 leader sequence a narrower mannosylation pattern is observed on the secreted and enriched GH-6 (Figure 5C and D) resulting in 24 – 30 hexoses (Figure 4B) as opposed to 29 – 37 hexoses (21). This implies that the unidentified translocon utilized by Celf_2022 may exhibit a modulating effect or strong relationship between mannosylation and secretion, whether it be a form of quality control or a greater degree of interaction with GT-39s. In addition, the conformation of the mannoprotein being exported cannot yet be ruled out as we have yet to identify other actinobacterial mannoproteins like Celf_2022 that are only POM targets when exported by their native translocons.

The lack of Celf_3184 secretion in the ΔCg_1014 strain initially implied some elements of the TAT translocon require mannosylation for functionality, however, the production of Celf_2022 with a TAT leader discredited this hypothesis (Figure S6C). While some components of both the SEC and TAT secretion pathways have been reported to be glycosylated in *M. tuberculosis* – with glycans composed of more than just mannose (77) – mannosylation does not appear to influence the functionality of either translocon in *C. glutamicum*. It is more likely that the abolishment of mannosylation in *C. glutamicum* results in the loss of functionality of one or more chaperones – a modification also observed on chaperones from *M. tuberculosis* (77) – specifically required for recombinantly produced Celf_3184 to be folded correctly, leading to its rapid degradation (89). Currently, the translocon utilized by Celf_2022 is unknown and to truly dissect its relationship with POM it must first be identified. Of note is the recently identified class of non-classically secreted proteins lacking a traditionally identifiable *N*-terminal signal peptide. Currently, it cannot be entirely ruled out that Celf_2022 may be a member of this class of proteins. While the study of non-classical secretion in bacteria is steadily growing – with reports in *Bacillus*, *Listeria*, *Staphylococcus*, *Streptococcus*, *Mycobacterium*, and others (63, 90–94) – the relationship between this novel secretion system and protein glycosylation in bacteria is under-reported.

### Conclusion

While POM has been known to occur in various actinobacterial species, a significant lack of information still exists about how this modification benefits them outside of the context of virulence. Even though they are closely related, two of the actinobacterial GT-39s investigated here showed very different substrate specificities for – and activities on – known actinobacterial mannoprotein targets. Moreover, the mannosylation deficient strain of *C. glutamicum* showed an increased susceptibility to tetracycline, chloramphenicol, and tobramycin suggesting POM may affect cellular permeability in some manner. Most importantly, is further evidence that POM does not only occur in a solely SEC dependent manner in actinobacteria, as proteins can be exported and mannosylated via the TAT pathway. Moreover, here we have presented evidence for a currently unidentified secretion pathway in actinobacteria that can mannosylate and export proteins without using either the SEC or TAT translocons with different levels and distributions of the modification based on the pathway used. These findings showcase how poorly understood actinobacterial POM is, both in how it is performed and what impact it has on the organisms that are capable of it.

Currently, *C. glutamicum* is a naïve recombinant host with the success of high-yield recombinant production occurring on a protein-to-protein basis for a variety of reasons. However, a recently developed molecular chaperone system has been shown to improve consistent recombinant protein production in the organism (89); a similar methodology will be utilized to facilitate the production and detection of heterologous GT-39s. While heterologous protein production and secretion in *C. glutamicum* benefits from the organism’s low abundance of secreted proteases (95) intracellular proteases can dramatically affect the yields of heterologous proteins (96), justifying the need for additional engineering of this strain.

## Methods

### Media, Strains, and Expression Conditions

All strains were grown in 2YT media (16 g/L tryptone, 10 g/L yeast extract, 5 g/L NaCl, BioShop Canada). NEB® Stable *E. coli* (NEB) was used for routine cloning and plasmid production. BL21 (DE3) *E. coli* and *C. glutamicum* ATCC13032 were used for recombinant protein production. Electrocompetent *C. glutamicum* were cultured in MBGT media (16 g/L tryptone, 10 g/L yeast extract, 5 g/L NaCl, 35 g/L glycine, 0.1% Tween-80) and outgrowths were performed in 2YT + 91 g/L sorbitol.

Single colonies of *C. glutamicum* expression constructs were inoculated into 25 mL 2YT containing 50 µg/mL kanamycin and 25 µg/mL nalidixic acid, then incubated overnight at 30°C with shaking at 180 RPM. The following day, overnight cultures were diluted into 250 mL 2YT expression cultures (containing 50 µg/mL kanamycin and 25 µg/mL nalidixic acid) to an OD_600_ ≈ 0.1 and incubated at 30°C, 180 RPM. Cultures were induced with 0.5 mM IPTG once they reached an OD_600_ ≈ 0.6 and allowed to express overnight at 30°C. Following induction, cultures were harvested by centrifugation at 5,000 × *g* for 10 mins at 4°C.

### C. glutamicum GT-39 Knockout Generation and Characterization

The gene encoding the sole GT-39 in *C. glutamicum* ATCC 13032 (Cg_1014) was replaced with a truncated and inactive mutant via homologous recombination assisted by the pK18mobsacB suicide vector. A synthetic gene composed of the truncated Cg_1014 gene containing only the cytoplasmic N-terminal domain (1 – 35 aa), the first transmembrane region (36 – 58 aa), and the extracellular C-terminal domain (506 – 520 aa) flanked by upstream and downstream regions of 1001 bps was synthesized (IDT), restriction cloned into pK18mobsacB using XbaI and SalI, and subsequently cloned into electrocompetent NEB® Stable *E. coli* (NEB). The sequenced knockout construct was transformed directly into electrocompetent *C. glutamicum* ATCC 13032 and selection for gene replacement was carried out using the previously established *sac*B methodology (62) with counterselection in the presence of 20% sucrose. Replacement of Cg_1014 with the truncated gene was confirmed via PCR with specific flanking primers (Table S2).

Growth curves were performed in triplicate in 2YT media (containing 50 µg/mL kanamycin and 25 µg/mL nalidixic acid) at 30°C and 180 RPM throughout and OD_600_ was measured spectrophotometrically. Antibiotic susceptibility was assayed with the Kirby-Bauer (KB) methodology at 30°C using the following antibiotic discs in quadruplicate: bacitracin (BAC) 10 U, vancomycin (VAN) 30 µg, cefotaxime (CTX) 30 µg, oxacillin (OXA) 1 µg, colistin (COL) 10 µg, tetracycline (TET) 30 µg, streptomycin (STR) 10 µg, novobiocin (NOV) 30 µg, chloramphenicol (CHL) 30 µg, tobramycin (TOB) 10 µg, kanamycin (KAN) 30 µg, ciprofloxacin (CIP) 5 µg, gentamicin (GEN) 10 µg, nalidixic acid (NAL) 30 µg, rifampicin (RIF) 5 µg, neomycin (NEO) 30 µg, and erythromycin (ERY) 15 µg (Fisher).

### Vector and Mannosylation Operon Design and Construction

The *E. coli/C. glutamicum* shuttle vector pTGR-5 (97) was received as a generous gift from Dr. Pablo Ravasi. To generate the high-level expression plasmid, pCGE-31, the Ptac region in pTGR-5 (XbaI – NheI fragment) was replaced with the lac UV5 + tandem Plac system from the expression vector pCW (98) utilizing synthetic primers (Table S2) and maintaining the *sod* RBS and spacing already present in pTGR-5. The improved expression levels of pCGE-31 are shown in Figure S9.

GT-39 genes from *C. glutamicum* ATCC 13032, *C. fimi* ATCC 484, and *C. flavigena* ATCC 482 were amplified from genomic DNA using specific primers containing NdeI (5’) and HindIII (3’) restriction sites (Table S2). The triple lac operator from pCW-MalET (98) was used to replace the single lac operator in pTGR-5 (97) using synthetic primers containing BamHI (5’) and NheI (3’) restriction sites (Table S2) while maintaining the RBS_sod_ and nucleotide spacing, generating the pCGE-31 shuttle vector (Figure S10A) for the recombinant expression of actinobacterial GT-39s in *C. glutamicum*. A synthetic operon used to assay POM *in vivo* using a known mannoprotein (Figure S10B) was designed and inserted upstream of the rrnB T1 terminator using NdeI (5’) and AvrII (3’) containing actinobacterial GT-39s followed by a single lac operator, RBS_sod_, and actinobacterial mannoprotein Celf_3184. Constructs were confirmed via restriction digest and sequencing, transformation of constructs into *C. glutamicum* was confirmed via plasmid rescue into NEB® Stable *E. coli* (NEB).

### Electrocompetent C. glutamicum

A single colony of *C. glutamicum* ATCC13032 or ΔCg_1014 was inoculated into 50 mL 2YT containing 25 µg/mL nalidixic acid and incubated overnight at 30°C with shaking at 180 RPM. The following day, 1 L MBGT containing 25 µg/mL nalidixic acid was inoculated to an OD_600_ ≈ 0.1 using the overnight culture and incubated at 30°C, 180 RPM. When the OD_600_ ≈ 0.25 – 0.25 (about 2 hours) 0.5 µg/mL ampicillin was added and the culture was allowed to continue incubating at 30°C, 180 RPM for an additional 1.5 hours.

Following incubation, cells were harvested at 5,000 × *g* for 10 mins at 4°C. Cells were resuspended in 150 mL 10% glycerol and centrifuged at 5,000 × *g* for 10 mins at 4°C a total of 3 times. Final cell pellets were resuspended in 10% glycerol to a final OD_600_ ≈ 200 and aliquots of 100 µL were stored at -80°C.

Electrocompetent *C. glutamicum* ATCC 13032 and ΔCg_1014 were transformed with an adapted protocol (99, 100). Competent cell aliquots were thawed on ice and allowed to incubate with 750 ng DNA for 10 mins prior to transformation. Cells and DNA were electroporated in 0.2 cm cuvettes using a BioRad Gene Pulser Mini at 2.5 kV for 4.80 – 5.20 ms. Immediately following, 1 mL 2YT + 91 g/L sorbitol was added to cells and outgrowths were placed at 46°C for 6 mins to inactivate the host restriction system and increase transformation efficiency. Cells were allowed to recover at 30°C, 180 RPM for 2 hours. Cells were harvested via centrifugation at 5,000 × *g* for 1 min and resuspended in 200 µL fresh outgrowth medium prior to plating on agar containing 50 µg/mL kanamycin and 25 µg/mL nalidixic acid. Plates were incubated at 30°C for 48 – 72 hours until colonies appeared.

All constructs in *C. glutamicum* were confirmed via plasmid rescue in *E. coli*. Miniprepped plasmid DNA from *C. glutamicum* constructs was transformed into electrocompetent *E. coli* for propagation, then confirmed by restriction digest and sequencing.

### Purification of Secreted HIS^6^-tagged Proteins

Cell pellets were harvested by centrifugation at 5,000 × *g* for 10 mins at 4°C and stored at -20°C. The spent culture media was clarified via centrifugation at 20,000 × *g* for 30 mins at 4°C and particulates were removed with a 0.45 µm PES bottle top filter (supp). A 10X stock of HisTrap A Buffer (1 M HEPES, 3 M NaCl, pH 8.0) was diluted to 1X with the clarified spent media, bringing the mixture to a final concentration of 100 mM HEPES, 300 mM NaCl, pH 8.0 prior to affinity chromatography.

All recombinantly produced proteins were enriched via affinity chromatography using Roche cOmplete™ His-Tag Purification Resin (Millipore-Sigma) and chromatography on an AKTA Start (Cytiva). Clarified spent media were loaded onto an equilibrated XK-16 column (Cytiva) containing 15 mL cOmplete™ resin with HisTrap A buffer at a flowrate of 2 mL/min. The column was washed with 3 CV of HisTrap A buffer prior to elution along a linear gradient (0 – 100%) of HisTrap B (100 mM HEPES, 300 mM NaCl, 500 mM imidazole, pH 8.0) over 5 CV. Fractions containing recombinant proteins were pooled, concentrated, and buffer exchanged into 50 mM HEPES, 150 mM NaCl, pH 7.4 using 20 mL VivaSpin concentrators with 10,000 MWCO (Cytiva).

### Isolation of C. glutamicum Membrane Proteins

Frozen cell pellets were resuspended (1 g/ 10 mL) in ConA buffer A (20 mM Tris, 0.5 M NaCl, 1.0 mM CaCl2 and 1.0 mM MnCl_2_, pH 7.4) with Sigma P-2714 protease inhibitor cocktail. The cells were lysed using an Emulsiflex-C5 (Avestin) at ≥ 20,000 psi. Following centrifugation at 6, 000 × g for 5 min at 4°C to remove debris, the supernatant was centrifuged again at 20,000 × g for 20 min at 4°C. The 20,000 × g pellet fraction was resuspended in 0.5 mL of ConA buffer A with 0.1% Triton X-100 and incubated at room temperature, using a tube roller, overnight at 4°C. Solubilized membranes were centrifuged at 100, 000 × g for 1 h at 4°C and the resultant supernatants were used for Western and lectin blotting.

### Western and Lectin Blot Protocol

Lectin blots were performed as described in the manufacturers data sheets. Briefly, the proteins of interest were separated on a 12% SDS page gel using a miniProtean system (Bio-Rad), which was then rinsed three times for 5 minutes each in excess Tris buffered saline pH 7.6 (TBS, 50 mM Tris, 150 mM NaCl, pH 7.6) before being blotted to PVDF membrane using a Trans-Blot Turbo transfer system (Bio-Rad). The transfer was performed in 48 mM Tris, 39 mM glycine using 2.5 A, 25 V for 8 mins. The protein-bearing PVDF membrane was then rinsed three times for 5 minutes each, in TBS and blocked for one hour in 5% BSA in TBS at room temp. The blocked membrane was washed three times for 5 minutes in TBS at room temp, then incubated overnight at 4°C in 0.05% Tween 20; 1mM CaCl_2_; 1mM MgCl_2_; 1mM MnCl_2_; 0.5 µg/mL ConA-FITC conjugated lectin (Millipore-Sigma); and 1:10, 000 AlexaFluor 647 anti-HIS_6_ (Bio-Rad) in TBS. The membrane then underwent three 10-minute washes in TBS pH 7.6 at room temp and was visualized on a Bio-Rad ChemiDoc. Molecular weight markers were Bio-Rad All Blue standards.

### Intact mass LC-MS analysis of Celf-3184 Mannosylated by Actinobacterial GT-39s

Intact mass analysis was performed using an Ultimate 3000 (Dionex/Thermo Fisher Scientific) linked to an LTQ-Orbitrap XL hybrid mass spectrometer (Thermo Fisher Scientific). 5 µg of each protein was injected on to a 2.1 × 30 mm Poros R2 column (Thermo Fisher Scientific) and resolved using the following rapid gradient: hold at 20% mobile phase A for 3 minutes, 20% - 90% mobile phase B in 3 minutes, hold at 90% mobile phase B for 1 minute. Mobile phase A was 0.1% formic acid in ddH2O and mobile phase B was acetonitrile. The flow rate was 3 mL/min with 100 µL split to the electrospray ion source. Optimal peak shape was achieved by heating the column and mobile phase to 80°C. The mass spectrometer was tuned for small protein analysis using myoglobin and the resolution was set to 15,000. Mass spectra were acquired from m/z 400 to 2,000 in the Orbitrap at 1 scan per second. The spectra acquired across the protein peak were summed and deconvoluted using MaxEnt 1 (Waters).

## Acknowledgements

This work was partially funded by a Natural Sciences and Engineering Research Council grant to WW.

---

**Figure S1:**
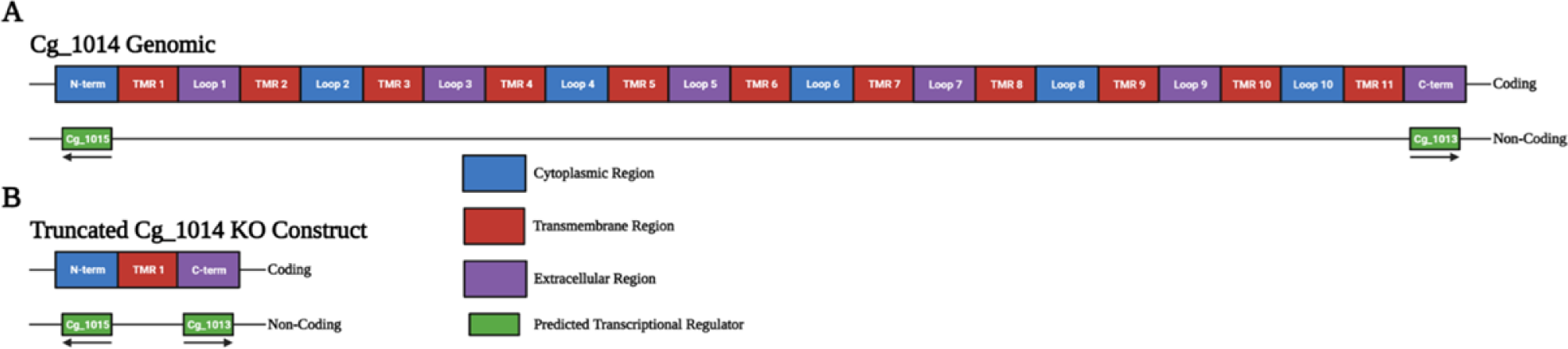
Schematic and predicted topology of genomic Cg_1014 and predicted transcriptional regulators on the non-coding strand (A) maintained in the inactivated knockout mutant (B). The *C. glutamicum* GT-39 Cg_1014 contains (as predicted by TMHMM-2.0) 11 transmembrane regions (TMR, red), 5 cytoplasmic loops (blue), and 6 extracellular loops (purple). The predicted cytoplasmic N-terminal and extracellular C-terminal domains also contain predicted transcriptional regulators for the neighbouring genes Cg_1013 and Cg_1015 (A). As catalytic activity is thought to be harboured on extracellular Loop 1 a truncated and inactive mutant was designed (B) containing only the N- and C-terminal domains connected by TMR 1, maintaining down- and upstream transcriptional effectors.

**Figure S2:**
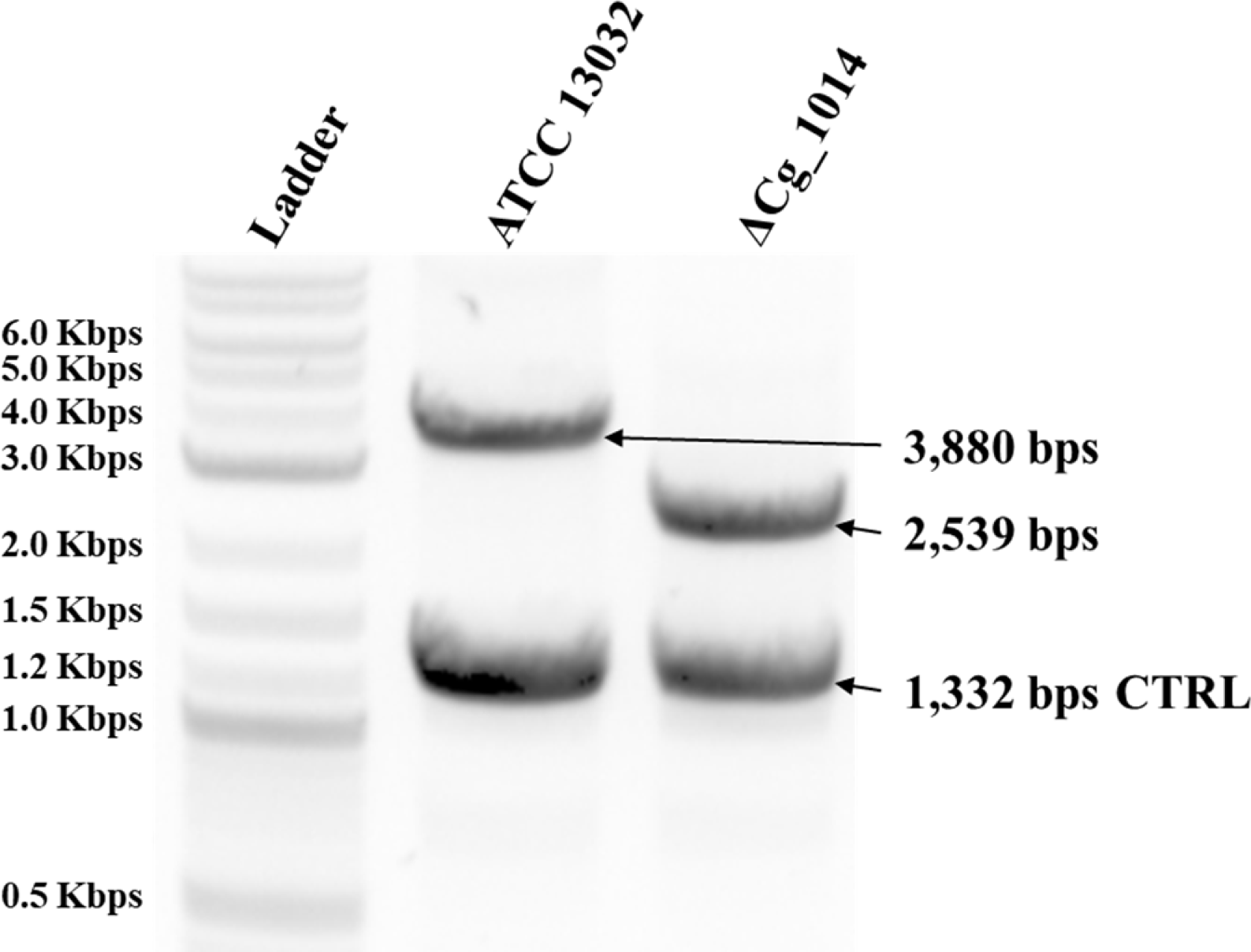
0.8% agarose gel showing genomic knockout of GT-39 in *C. glutamicum* ATCC 13032 and ΔCg_1014. Primers specific to the flanking regions ≈ 1,000 bps up- and downstream of Cg_1014 were used to identify successful homologous recombination events. In ATCC 13032 this amplicon is 3,880 bps and when Cg_1014 is replaced by the truncated inactive sequence the amplicon is 2,539 bps. This decrease exactly corresponds to the 1,341 bps removed from Cg_1014 (Δ59 – 505 aa) to generate the mutant gene. An additional amplicon of 1,332 was also produced using primers specific for the *C. glutamicum* homologue of the maltose binding protein (MBP) as a control.

**Figure S3:**
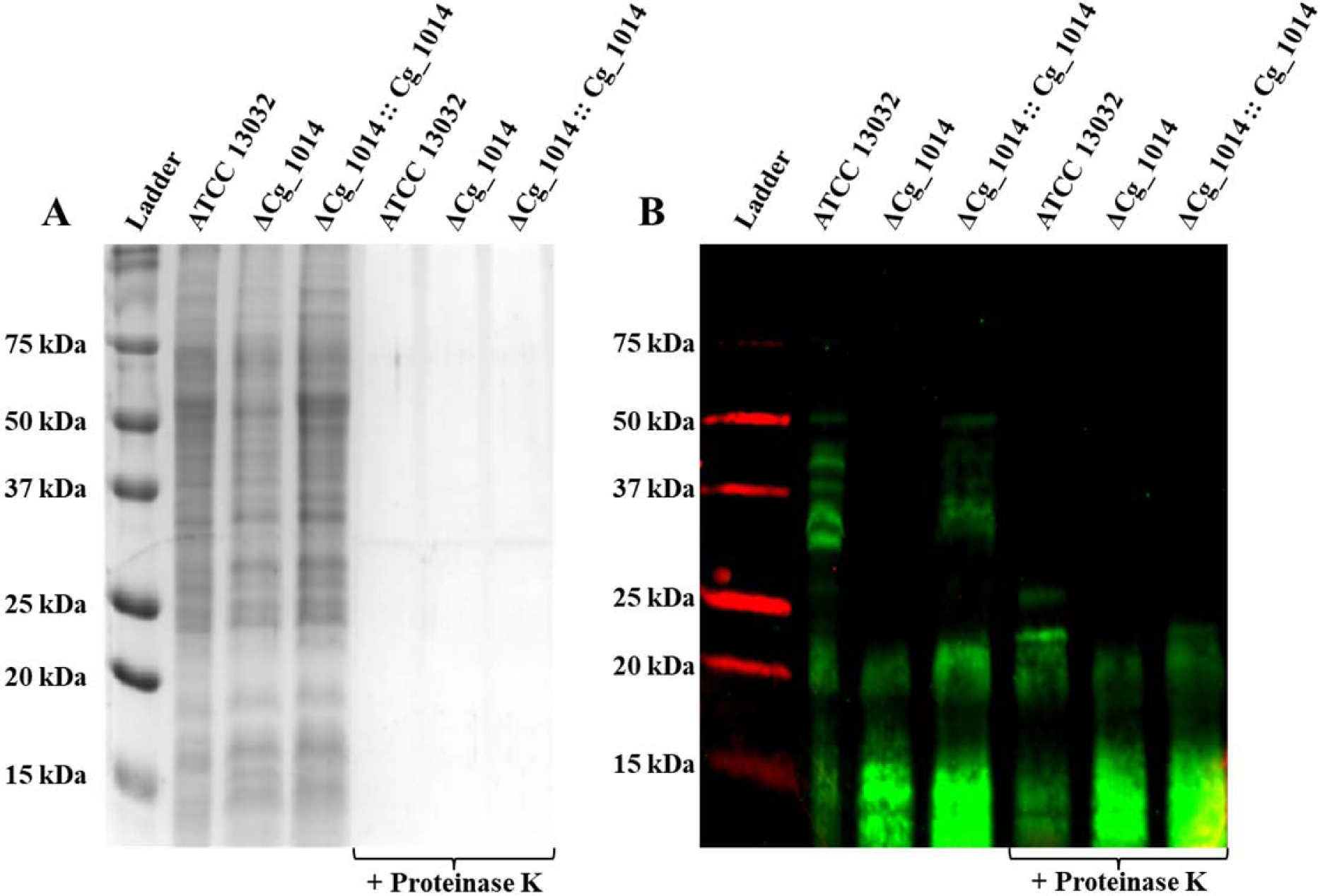
Coomassie stained 15% SDS-PAGE (A) and ConA-FITC (green) lectin blot (B) of *C. glutamicum* ATCC 13032, ΔCg_1014, and ΔCg_1014:Cg_1014 membrane fractions following digestion with proteinase K. Coomassie stained gel (A) shows membrane fractions before (Lanes 2 – 4) and after proteinase K digestion (Lanes 5 – 7). Digestion of proteins in each membrane fraction is evident by lack of Coomassie strained bands in proteinase K treated samples (A). ConA reactive smears in lectin blot (B) of membrane fractions from *C. glutamicum* ATCC 13032, ΔCg_1014, and complemented strains are attributed to the presence of LAM in the samples. Distinct bands that remain in ATCC 13032 and ΔCg_1014:Cg_1014 membrane fractions following proteinase K digest (Lanes 5-7) are attributed to protease resistant mannoproteins as POM is known to confer proteolytic resistance. Molecular weight standards are the Bio-Rad All Blue ladder.

**Figure S4:**
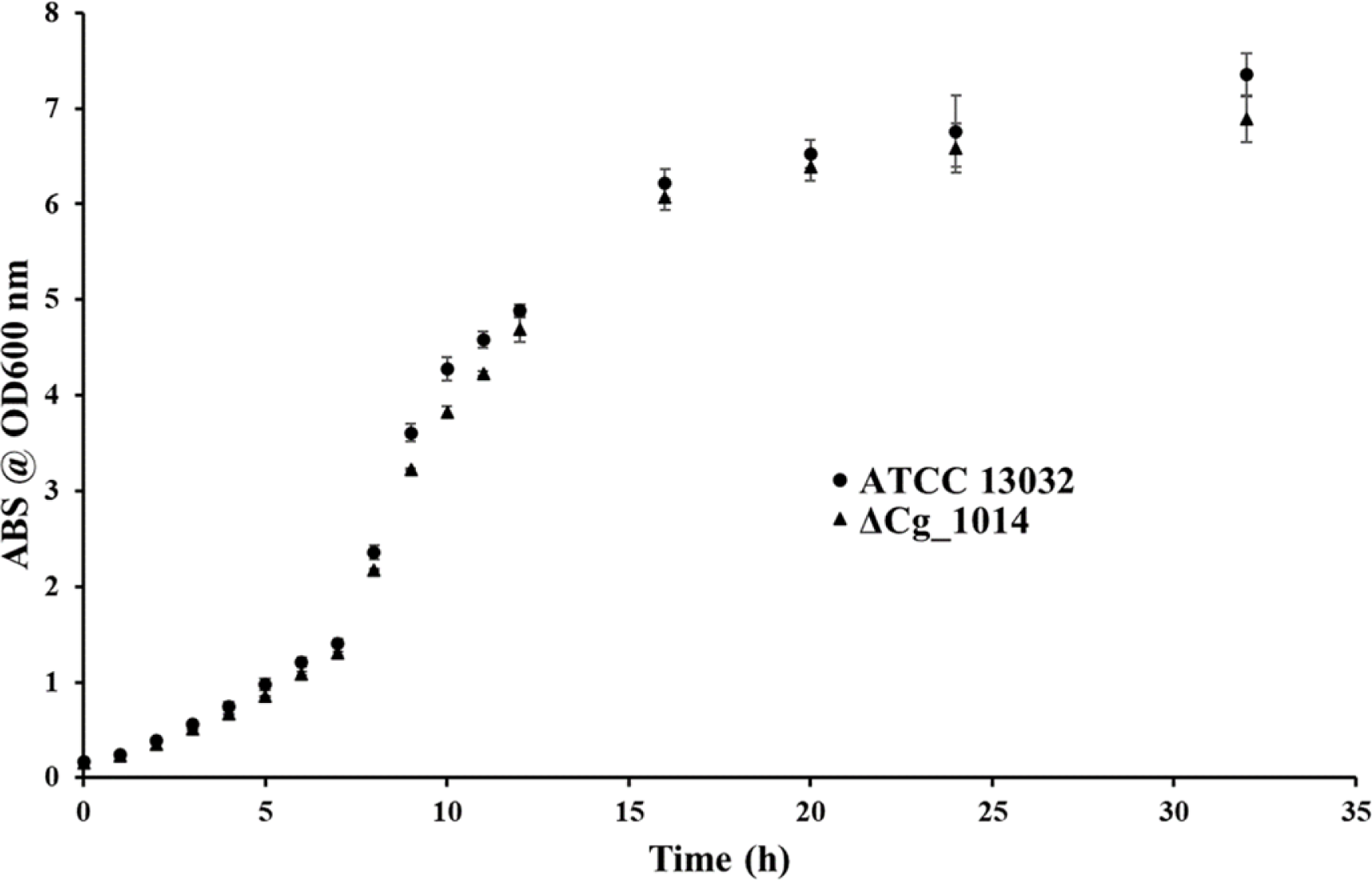
Effects of GT-39 inactivation on the growth of *C. glutamicum* ΔCg_1014. No significant differences to the ATCC 13032 strain were evident at any stage of growth in the ΔCg_1014 strain

**Figure S5:**
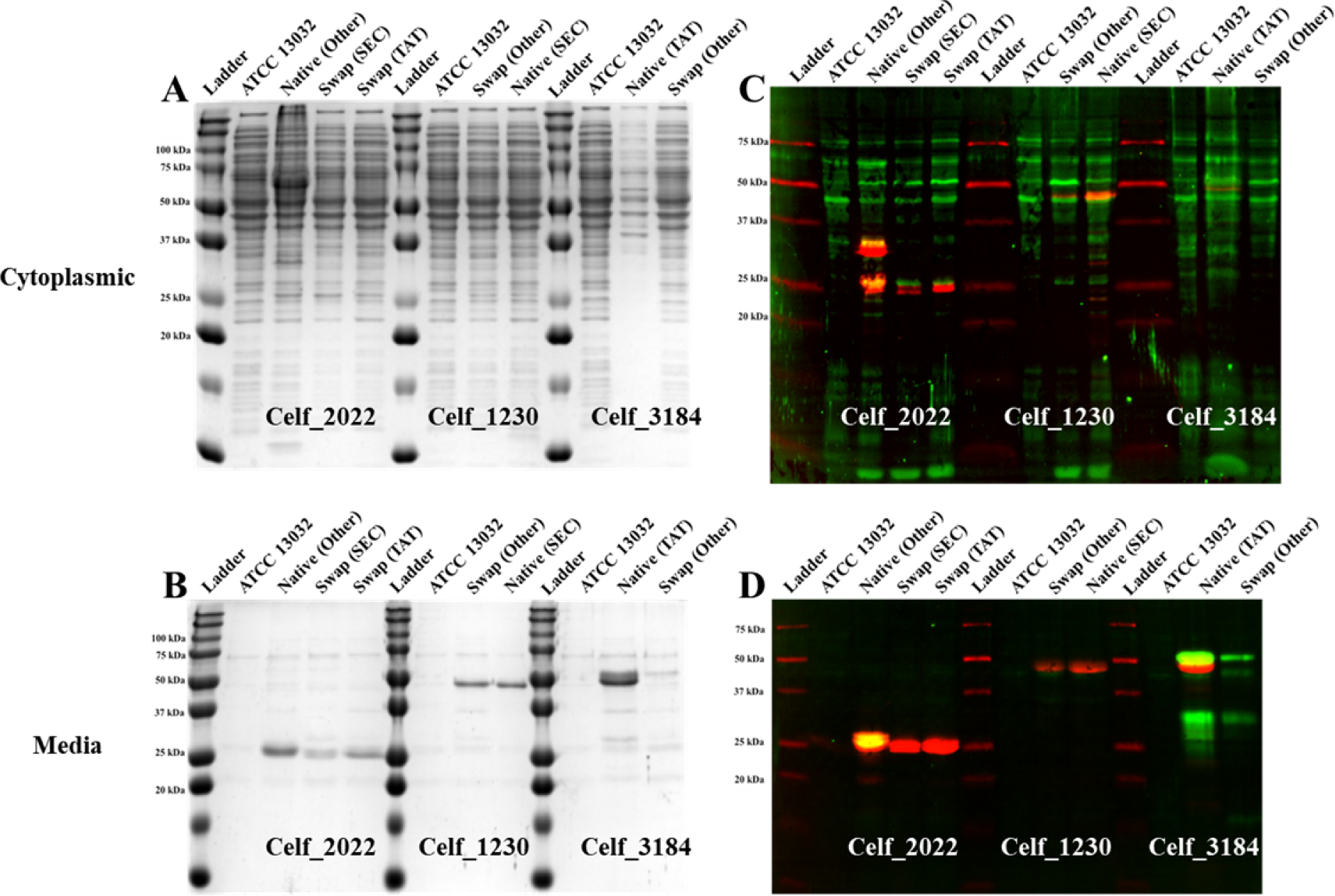
Coomassie stained 15% SDS-PAGE (A and C) and ConA-FITC (green) lectin blot (B and D) of *C. glutamicum* ATCC 13032 cytoplasmic (A and B) and undiluted spent culture media fractions (C and D) producing leader swap constructs. Celf_2022 was expressed with its native “Other” and swapped SEC, and TAT leaders, Celf_1230 was expressed with its native SEC and swapped “Other” leader, and Celf_3184 was expressed with its native TAT and swapped “Other” leader. Less overall protein is observed in the cytoplasm when Celf_2022 has its native “Other” leader swapped to either a SEC or TAT leader with no mannosylation evident on the secreted material, but secretion of the protein itself is not affected. Secreted Celf_1230 levels and its mannosylation status do not change when exchanging the native SEC leader to the “Other” leader, but cytoplasmic levels do decrease. Both secreted and cytoplasmic levels of Celf_3184 decrease when its native TAT leader is replaced by the “Other” leader, however the secreted material is much more homogenously mannosylated.

**Figure S6:**
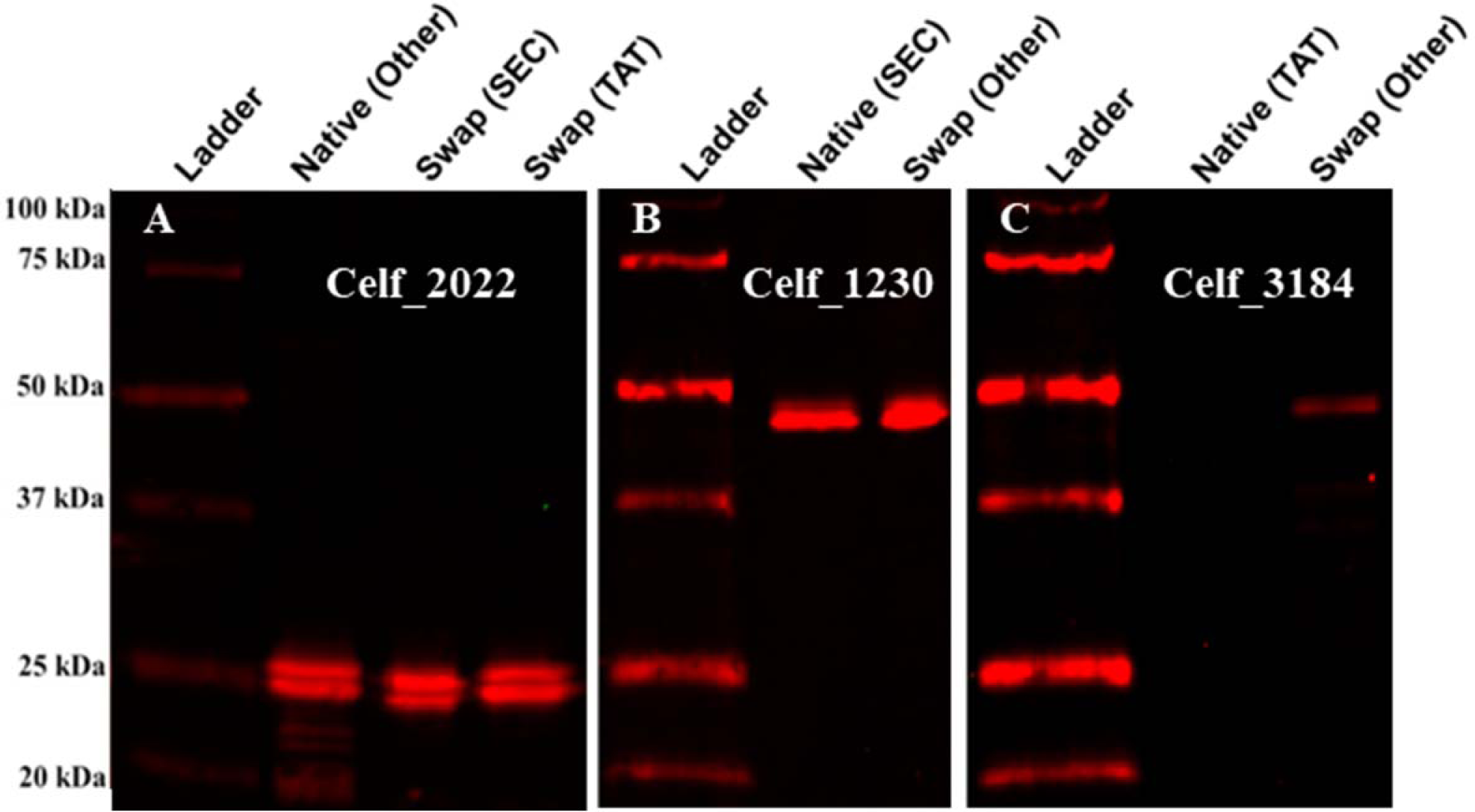
ConA-FITC (green) and Anti-HIS-Alexa647 (red) blot of spent culture media enriched recombinant Celf_2022 expressed with “other”, SEC, and TAT leaders (A), Celf_1230 expressed with SEC and “other” leaders (B), and Celf_3184 expressed with TAT and “other” leaders (C) produced in *C. glutamicum* ΔCg_1014. All proteins were produced in the POM deficient strain of *C. glutamicum*, indicated by the lack of ConA lectin reactivity (green). When lead by its native “other” leader sequence, Celf_2022 is secreted but not mannosylated in ΔCg_1014 (A). Secretion and mannosylation status are retained following the replacement of this “other” leader sequence by a SEC or TAT signal peptide. Replacement of the SEC signal peptide of Celf_1230 (a non-mannosylated protein) by the “other” leader sequence results in no change to secretion or mannosylation pattern (B). The native TAT led Celf_3184 is not produced or secreted in the ΔCg_1014 strain (C), but replacement of the TAT signal peptide by the “other” leader sequence results production and secretion of non-mannosylated Celf_3184. Molecular weight standards are the Bio-Rad All Blue ladder.

**Figure S7:**
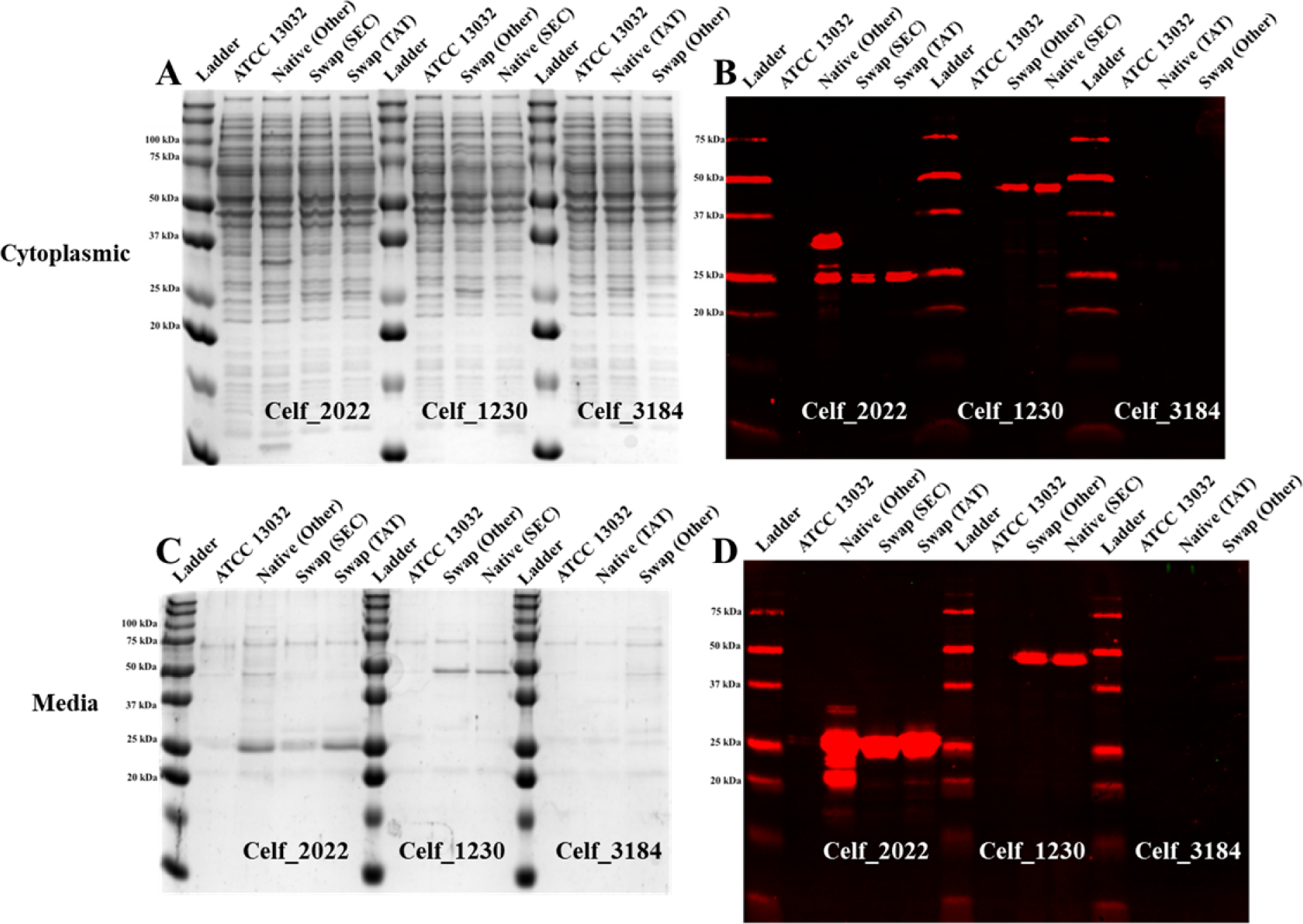
Coomassie stained 15% SDS-PAGE (A and C) and ConA-FITC (green) lectin blot (B and D) of ΔCg_1014 *C. glutamicum* cytoplasmic (A and B) and undiluted spent culture media fractions (C and D) producing leader swap constructs. Celf_2022 was expressed with its native “Other” and swapped SEC, and TAT leaders, Celf_1230 was expressed with its native SEC and swapped “Other” leader, and Celf_3184 was expressed with its native TAT and swapped “Other” leader. No mannosylation is evident on any of the constructs as they were expressed in the ΔCg_1014 mutant. Cytoplasmic and secreted protein abundance is very similar to these constructs expressed in the ATCC 13032 strain: however, Celf_3184 with its native TAT leader is not detectable (via Western blot and activity assay) in either the cytoplasmic or spent media fractions. Replacement of the TAT leader with the “Other” leader appears to restore the production and export of this protein.

**Figure S8:**
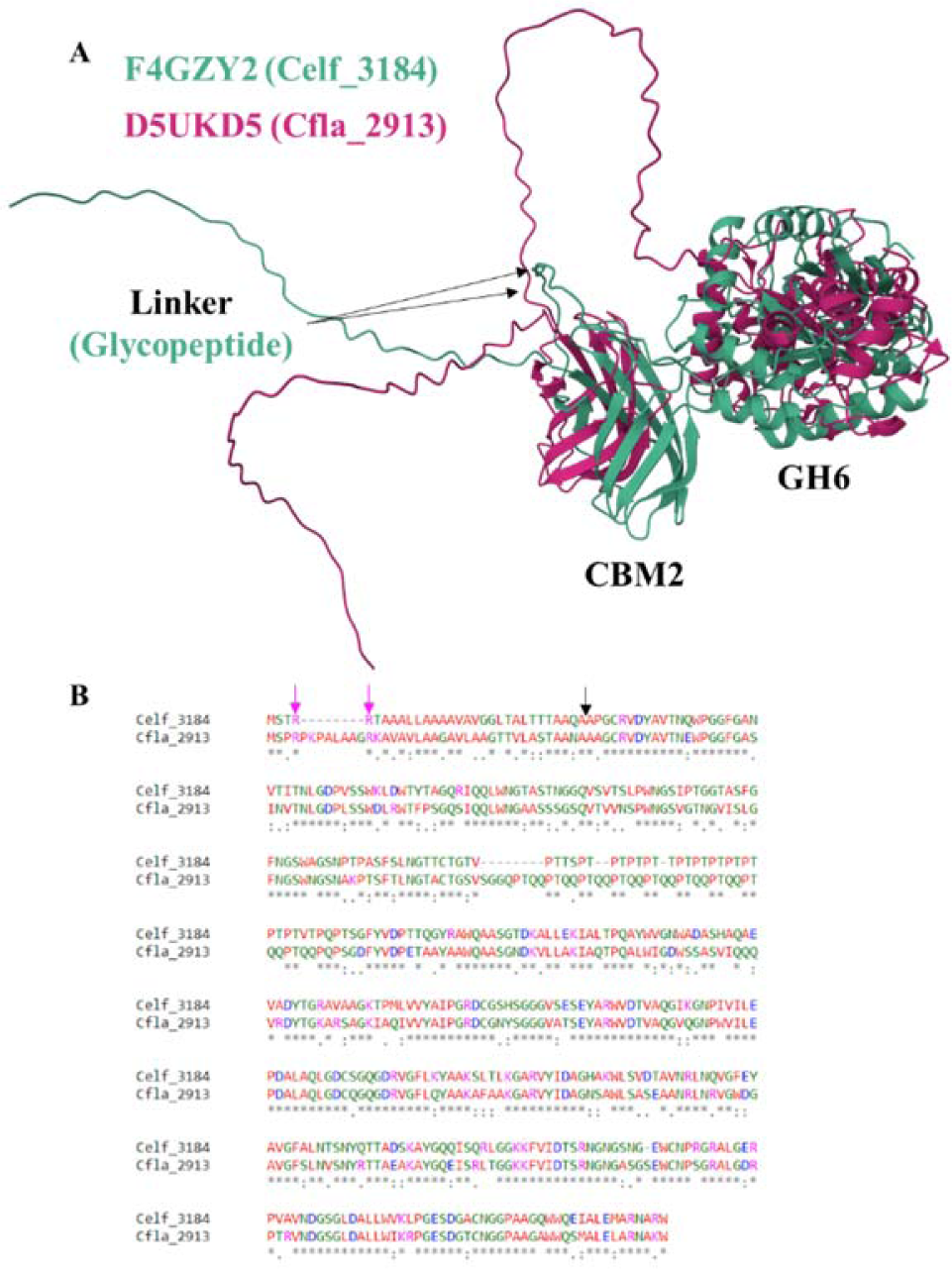
Alphafold predicted structures (A) comparing Celf_3184 (F4GZY2, green) and Clfa_2913 (D5UKD5, magenta) and MUSCLE 3.8 sequence alignment (B). Cfla_2913 – the *C. flavigena* homologue of the *C. fimi* GH6, Celf_3184 – contains a number of insertions in the linker region spanning the CBM2a and GH6 domains. In Celf_3184, this linker has been identified as the glycopeptide containing *O*-mannosyl glycans. The longer linker domain of Cfla_2913 promotes a significantly different orientation to this domain, potentially moving the glycosylation sites away from the active site of the *C. glutamicum* GT-39 (A). The *C. flavigena* and *C. glutamicum* GT-39s only share 40.5% global sequence homology. The twin arginine (RR) motif of Cfla_2913 contains an 8 amino acid insertion (B) – compared to the same motif in Celf_3184 – causing predictive tools like SignalP5.0 to misrepresent their identity. The full length Cfla_2913 protein also contains the hydrophobic and C-terminal regions indicative of TAT proteins. The C-terminal region often ends with a short (A-X-A) motif specifying cleavage by the signal peptidase. Predicted signal peptide cleavage of both proteins indicated by black arrow and twin arginine motif indicated by magenta arrows.

**Figure S9:**
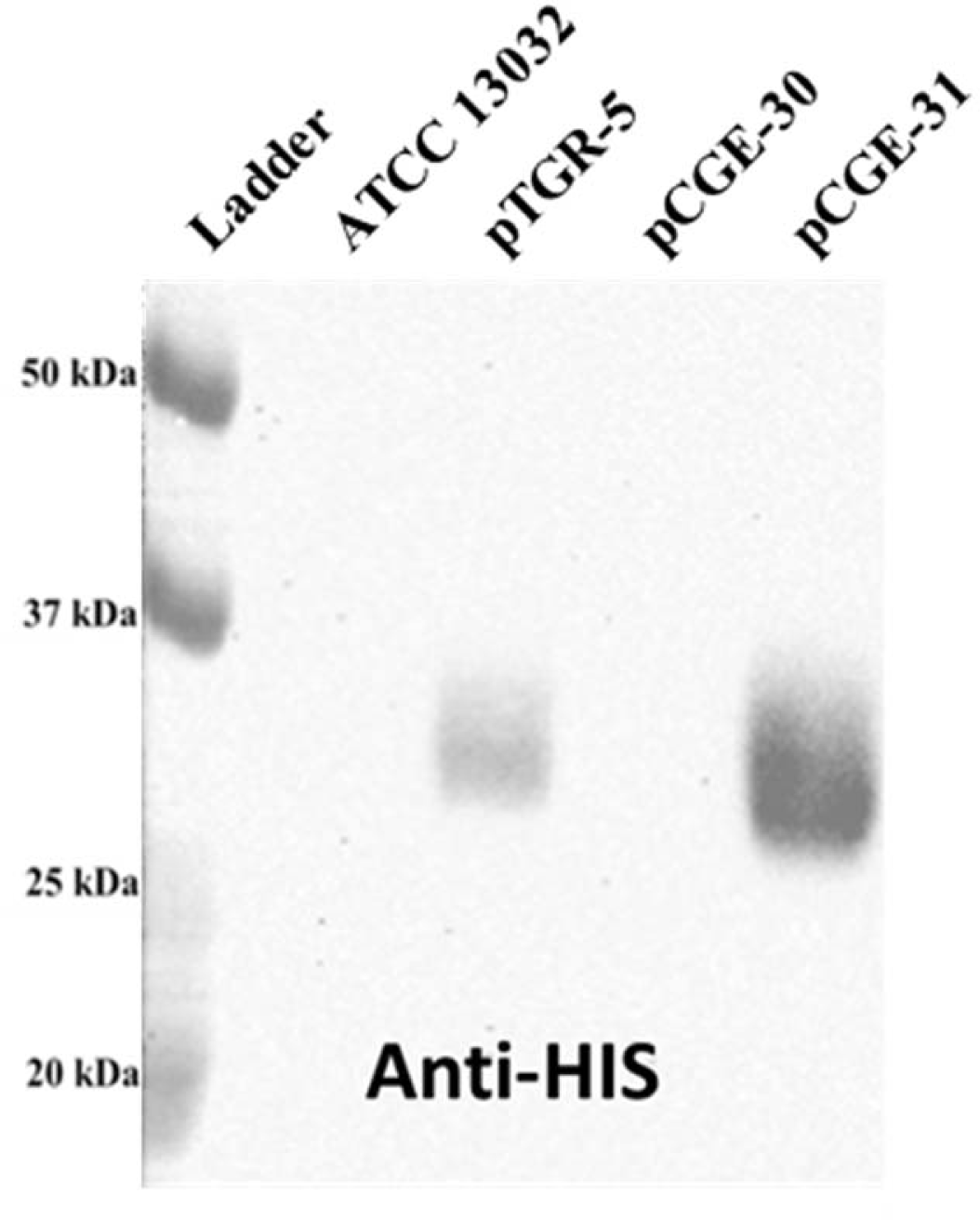
Anti-HIS-HRP western blot showing expression level differences of eGFP produced in *C. glutamicum* by pTGR-5 and pCGE-31. The *E. coli*/*C. glutamicum* shuttle vector pCGE-31 can drive higher expression levels of recombinant proteins in *C. glutamicum* than its parent pTGR-5. The inclusion of the sod RBS is critical to its functionality in *C. glutamicum*, as pCGE-30 uses the triple promoter system and RBS from pCW and is incapable of producing as much eGFP as either sod RBS containing vector.

**Figure S10:**
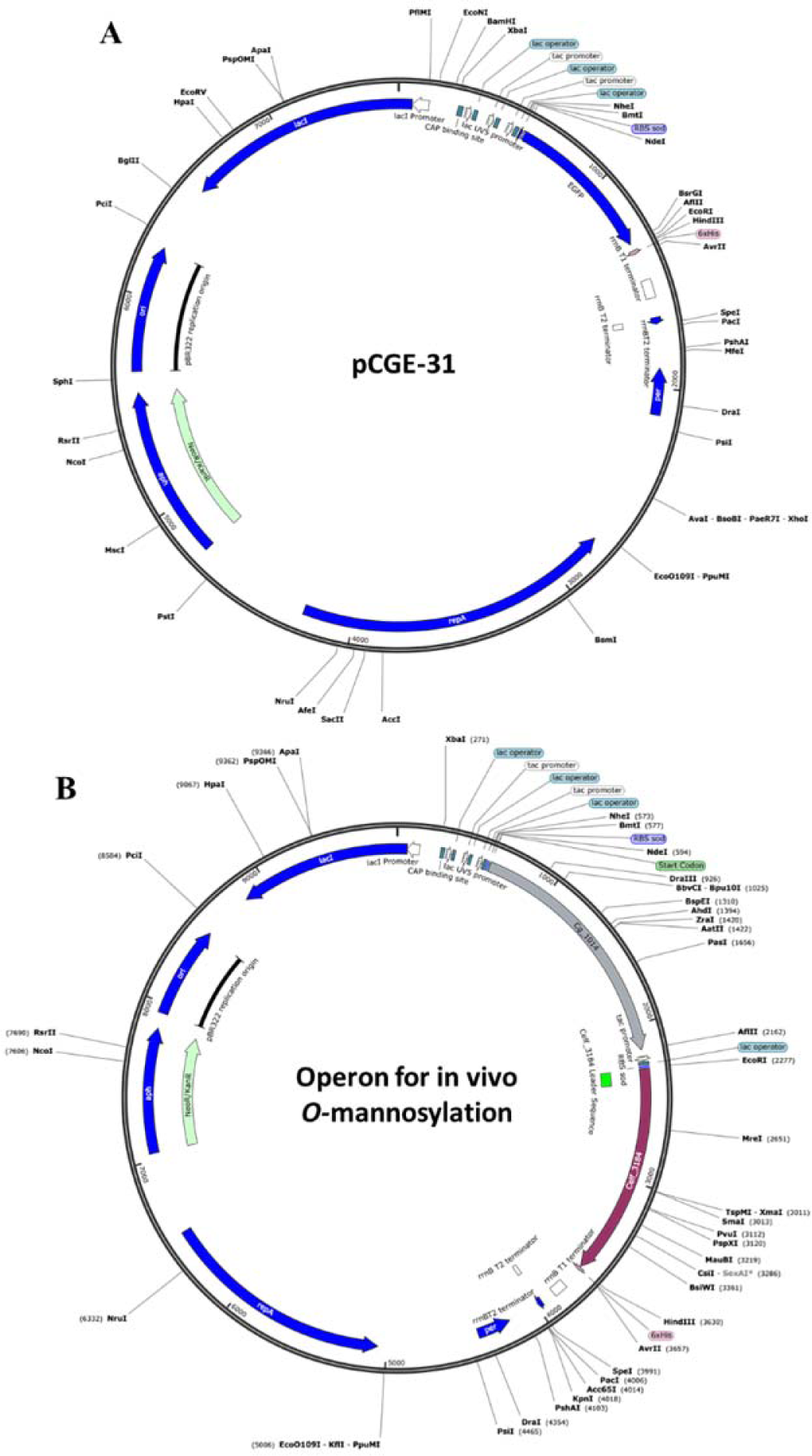
Plasmid maps of pCGE-31 used for recombinant expression of GT-39s (A) and *O*-mannosylation operons used for co-expression of GT-39s with target actinobacterial mannoproteins (B). Features of pCGE-31 *E. coli*/*C. glutamicum* shuttle vector (A). The pCGE-31 vector originates from pTGR-5 and received the triple promoter system from pCW via restriction cloning. Actinobacterial GT-39 genes amplified from genomic DNA replaced eGFP using NdeI and HindIII restriction sites. Features of the *O*-mannosylation operon constructs (B). A synthetic secondary ORF containing the Celf_3184 gene was added via restriction cloning (NdeI – AvrII) under the control of Ptac and also utilizing the sod RBS.

**Table S1:** Global sequence similarity of actinobacterial GT-39s compared to *S. cerevisiae* PMT1. Global sequence similarity was determined using EMBOSS Needle pairwise sequence alignment.

|  | *S. cerevisiae*<br>(PMT1) | *M. tuberculosis* | *M. smegmatis* | *C. glutamicum* | *C. fimi* | *C. flavigena* |
| --- | --- | --- | --- | --- | --- | --- |
| *S. cerevisiae* (PMT1) | 100.0% | 25.7% | 25.1% | 26.0% | 25.4% | 23.2% |
| *M. tuberculosis* | 25.7% | 100.0% | 83.9% | 55.9% | 46% | 46.9% |
| *M. smegmatis* | 25.1% | 83.9% | 100.0% | 57.7% | 46% | 48.7% |
| *C. glutamicum* | 26.0% | 55.9% | 57.7% | 100.0% | 42% | 40.5% |
| *C. fimi* | 25.4% | 46.0% | 46% | 42.0% | 100.0% | 50.5% |
| *C. flavigena* | 23.2% | 46.9% | 48.7% | 40.5% | 50.5% | 100.0% |

**Table S2:**
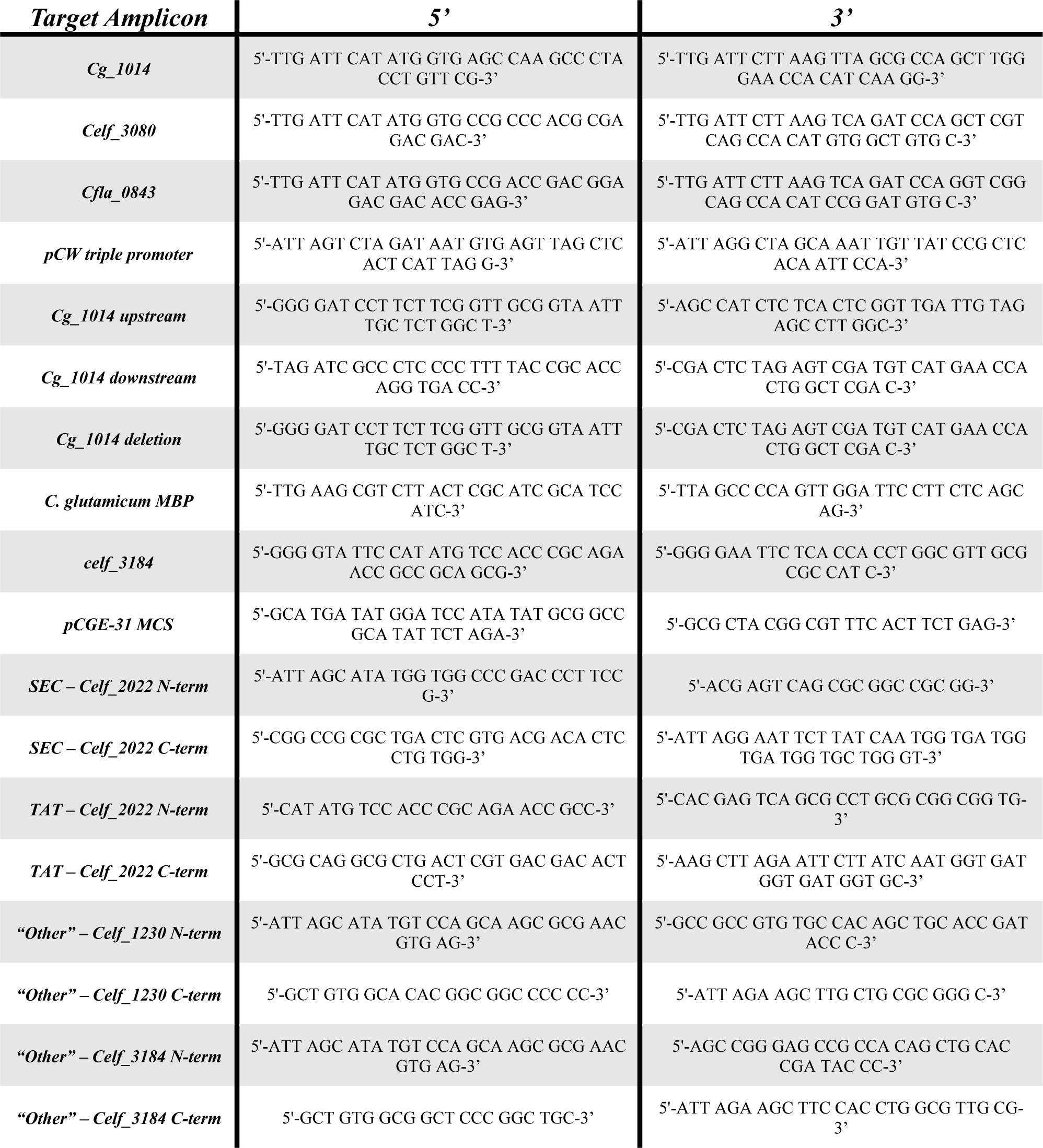
Selected primers used in this study.

